# Yeast Heterochromatin Only Stably Silences Weak Regulatory Elements by Altering Burst Duration

**DOI:** 10.1101/2023.10.05.561072

**Authors:** Kenneth Wu, Namrita Dhillon, Antone Bajor, Sara Abrahamson, Rohinton T. Kamakaka

## Abstract

The interplay between nucleosomes and transcription factors leads to programs of gene expression. Transcriptional silencing involves the generation of a chromatin state that represses transcription and is faithfully propagated through DNA replication and cell division. Using multiple reporter assays, including directly visualizing transcription in single cells, we investigated a diverse set of UAS enhancers and core promoters for their susceptibility to heterochromatic gene silencing. These results show that heterochromatin only stably silences weak and stress induced regulatory elements but is unable to stably repress housekeeping gene regulatory elements and the partial repression did not result in bistable expression states.

Permutation analysis of different UAS enhancers and core promoters indicate that both elements function together to determine the susceptibility of regulatory sequences to repression. Specific histone modifiers and chromatin remodellers function in an enhancer specific manner to aid these elements to resist repression suggesting that Sir proteins likely function in part by reducing nucleosome mobility. We also show that the strong housekeeping regulatory elements can be repressed if silencer bound Sir1 is increased, suggesting that Sir1 is a limiting component in silencing.

Together, our data suggest that the heterochromatic locus has been optimized to stably silence the weak mating type gene regulatory elements but not strong housekeeping gene regulatory sequences which could help explain why these genes are often found at the boundaries of silenced domains.

## Introduction

The DNA in a eukaryotic nucleus is wrapped around histones to form nucleosomes and non-histone proteins such as transcription factors interact with DNA and nucleosomes to form chromatin. The interplay between transcription factors and nucleosomal packaged DNA sequences ultimately leads to stable programs of gene expression that are critical for cell differentiation and proper development of organisms ^1^.

### Transcription Factors and Transcription Activation

Regulated transcription of genes requires sequence specific transcription activators as well as the general transcription factors. The latter bind sequences in the core promoter and mediate the formation of the preinitiation complex while sequence-specific transcription factors (TFs) bind to upstream activating sequence (UAS) enhancers to regulate transcription ^2,3^. Most yeast TFs are not essential for viability but are required for growth in stress conditions ^4^ although there are a handful of essential yeast TFs referred to as pioneer factors or general regulatory factors (GRF)”and Rap1, Abf1 and Reb1 are canonical GRFs ^5–9^.

Genes in yeast have been classified into distinct groups based on their enhancer/promoter architecture and transcription factor requirements. The largest class, numbering approximately 2500, are constitutively transcribed at very low levels typically generating between one and three mRNAs/hour ^10–12^. Approximately 1000 genes are considered stress responsive genes and are transcribed at very low levels in rich media but are highly induced under stress conditions ^11,13^ and are usually regulated by a single key transcription factor. The growth/housekeeping genes, refer to genes whose expression is highest during rapid growth and encode for ribosomal proteins and glycolytic enzymes. This class of genes is regulated by combinations of _the GRFs_^11,14-21^.

Nucleosome architecture is integral to gene regulation by transcription factors and the preinitiation complex cannot assemble at a core promoter if that element is packaged into a nucleosome ^22,23^. Genome-wide nucleosome profiling has revealed a high degree of nucleosome organization at the regulatory regions of most genes with well-positioned +1 and -1 nucleosomes flanking a nucleosome depleted region (NDR) of variable width and depth encompassing the UAS enhancer and core promoters ^8,24–28^. The removal and/or the maintenance of an NDR requires some transcription factors ^21,29,30^ as well as specific chromatin remodelling factors ^31,32^. RSC, SWI/SNF, INO80, and ISW2 act at the NDR to position the +1 and -1 nucleosomes. RSC and SWI/SNF push the +1 and -1 nucleosomes away from the NDR ^19,33,34^. In opposition to the pushing remodellers are INO80 and ISW2 remodellers which reposition the +1 and -1 nucleosomes towards the NDR ^34–36^.

In eukaryotes, transcription occurs in bursts due to the thermodynamics of sequence specific and general transcription factor binding and nucleosome mobility. Gene transcription can be divided into three steps. First, how frequently one observes transcription (transcription initiation frequency or burst frequency). This represents activator binding, nucleosome remodelling, preinitiation complex (PIC) formation and transcription initiation. Once transcription is initiated, multiple polymerases can be loaded during a transcription burst. Finally, the transcription burst is of a limited duration, and so the promoter switches from an active to a quiescent state ^37–41^. TF binding affects burst frequency ^42^ while TF dwell time determines burst duration which in turn depends on the affinity of the factor for the chromatinized binding site suggesting that the burst duration is encoded in the sequence ^43–53^. Nucleosomes play an important role in bursting, since they influence accessibility of sequences and the dwell time of transcriptional activators bound to DNA ^54^.

### Sir Proteins and Silencing

The yang of the yin of gene activation is the gene repression machinery. Transcriptional gene silencing is a form of gene repression that leads to the stable inactivation of genes. It involves the generation of a chromatin state that inhibits transcription. Silencing is mediated by DNA sequence elements called silencers and at the *HMR* locus, the *HMR-E* and *HMR-I* silencers flank the silenced *MATa1* and *MATa2* genes. *HMR-E* is bound by ORC, Rap1 and Abf1 proteins while *HMR-I* is bound by Abf1 and ORC^55,56^. The *HMR-E* bound ORC proteins aid in the binding of Sir 1 to the *HMR-E* silencer ^57–59^. Sir1 along with the other silencer bound proteins help recruit Sir2, Sir3 and Sir4. The Sir2–4 complex is responsible for the deacetylation of lysine16 in histone H4 in nucleosomes to promote further binding of the Sir complex to unacetylated nucleosomes ^56^. Sir protein bound chromatin is less accessible to molecular probes suggesting that these proteins function in gene repression by steric hindrance of transcription activators or general transcription factors ^60–63^ although some studies suggest that silencing occurs post-recruitment of these factors ^64,65^.

The transcriptionally silent state is highly stable and silencing of the *MAT* genes is rarely lost ^66^. In cells with mutations in either Sir1 ^67^ or the silencers ^68^, the fidelity of silencing is reduced, and two populations of cells are observed, some where the genes are fully silent and some where the genes are fully active. Furthermore, these expression states, once established, are propagated across several cell generations leading to two metastable populations of cells resulting in bistability of expression states ^67–69^.

Classical studies on silencing have focused on a set of UAS enhancer and core promoters of stress inducible or mating type genes that are controlled by single transcription activators ^55,56^. Housekeeping genes, constitute around a third of the genes in yeast, but the ability of these genes to be silenced has not been investigated and the functional relationships and communications between the activation machinery located at UAS enhancers and promoters and the silencing machinery recruited by the silencers is unclear.

In this manuscript we analysed silencing of a set of UAS enhancers and core promoters of varying strengths. We chose different classes of housekeeping, stress induced as well as well as weak constitutively expressing genes. The aim was to characterize the extent and form of silencing using multiple reporter systems. We show that the wild type silencers weakly affect highly transcribed housekeeping genes and partially repressed these genes by altering the burst duration. While partial repression of the housekeeping gene elements was observed with the native silencers, this repression was not stably propagated and did not result in bistable expression states. While the native silencers were unable to effectively repress housekeeping genes, analysis with synthetic silencers showed a correlation between the amount of Sir1 bound to a silencer and repression of these strong enhancer/promoters.

Analysis of permutations of UAS enhancers and core promoters demonstrated that sequences in both the enhancer and the core promoter determine the extent to which a regulatory sequence is susceptible to gene repression. Mutant analysis highlighted a role for chromatin remodellers in preventing an enhancer or promoter from being repressed suggesting that inhibiting nucleosome mobility may be an important mechanism by which the Sir proteins mediate silencing. Our data suggest that the silenced locus has been optimized for just enough silencing to stably silence the mating type gene UAS enhancers and core promoters but not for the silencing of strong activating regulatory sequences.

## Materials and Methods

Yeast strain genotypes can be found in Table 1.

**Table 1.**
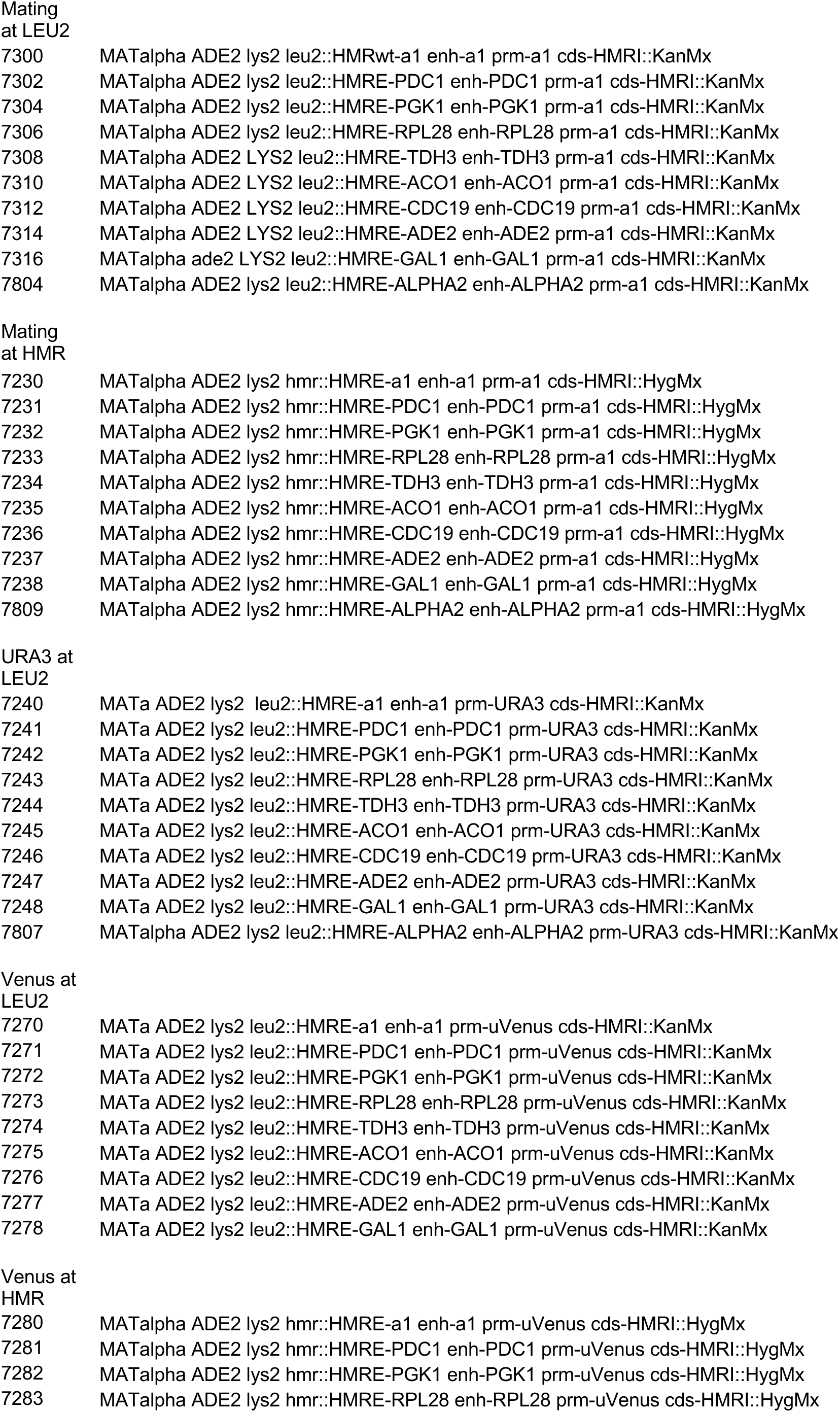
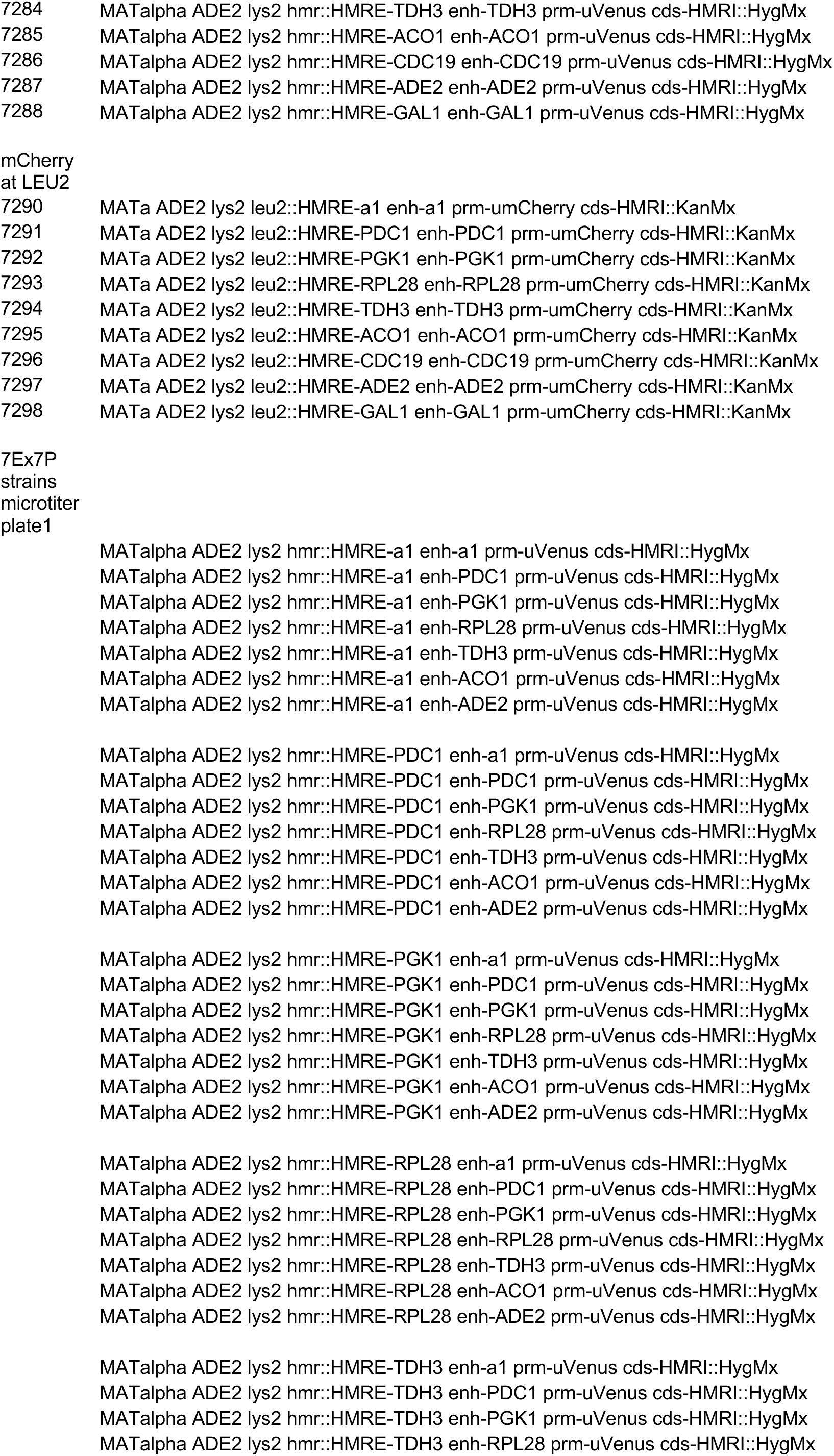
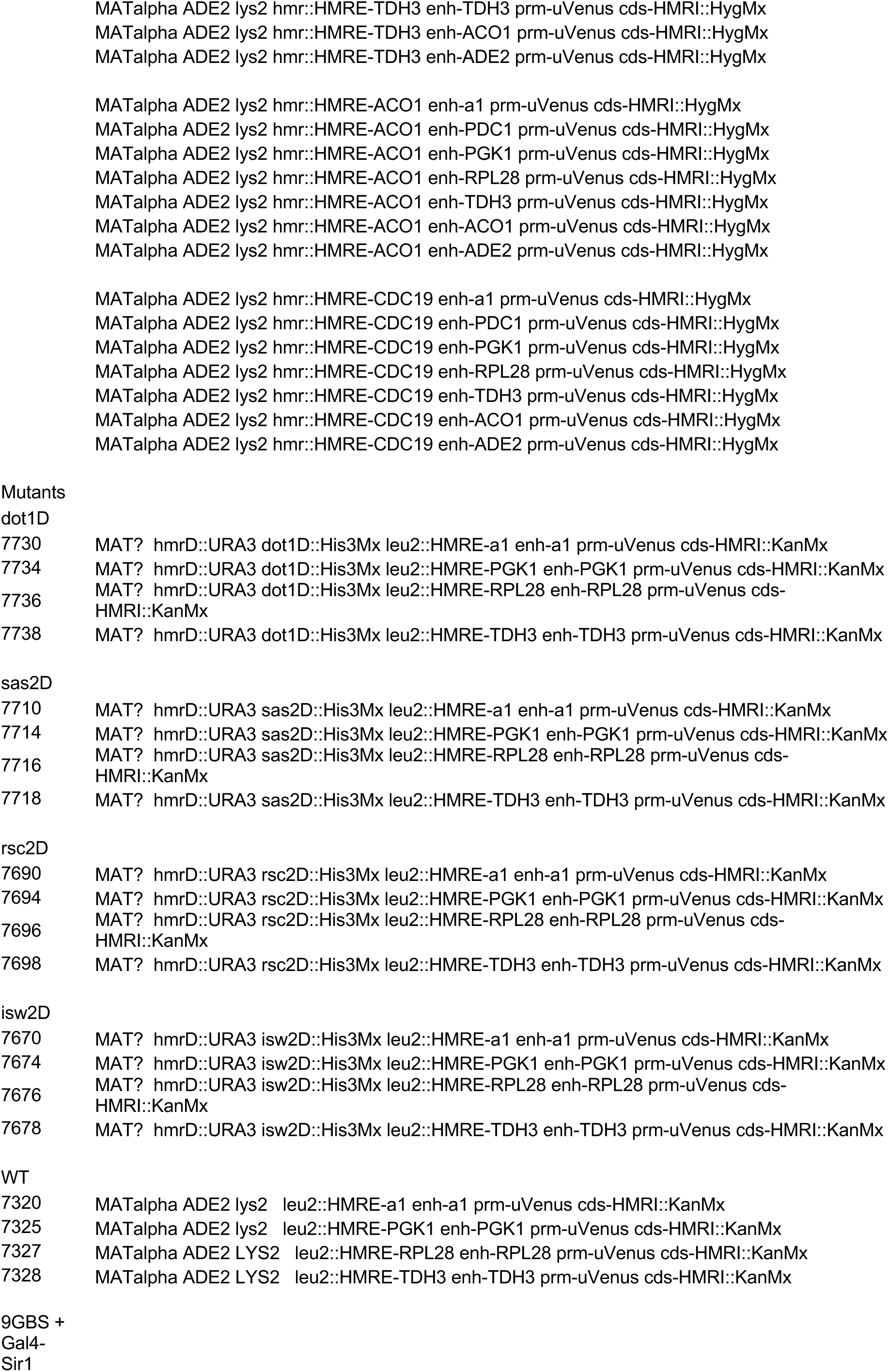
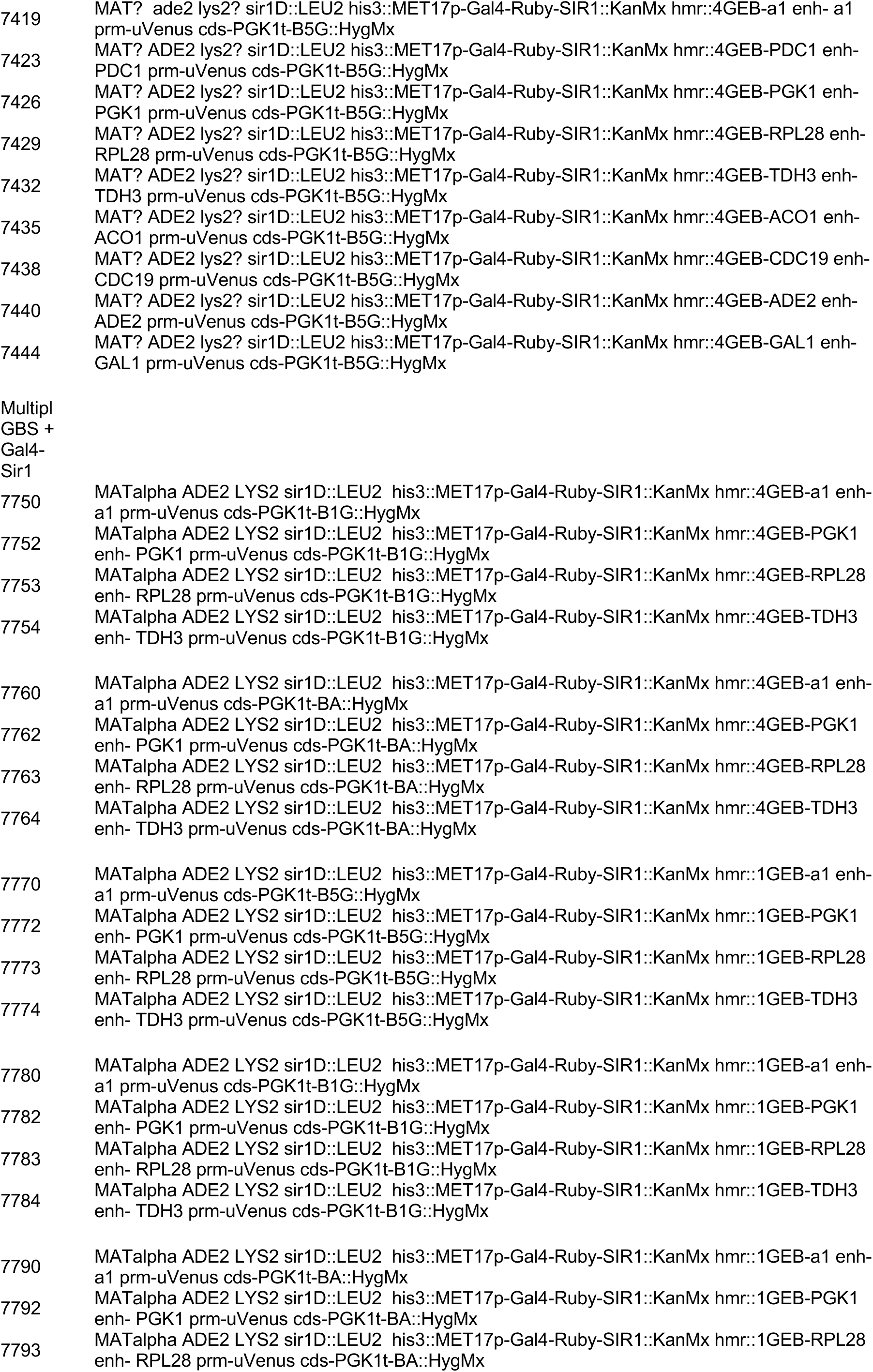
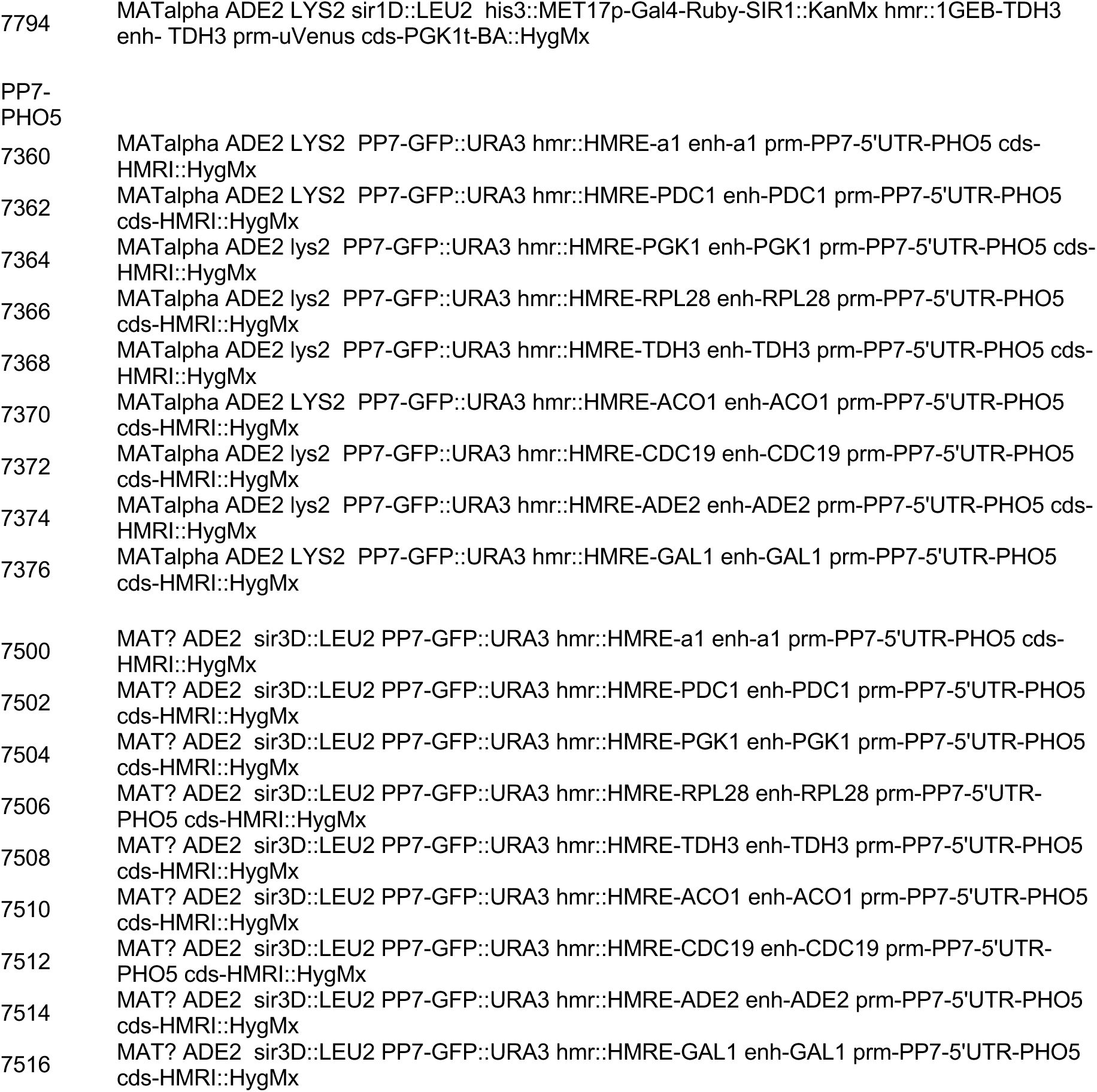
Genotypes of Yeast strains used in this study.

### Semi-Quantitative mating assays

Semi-quantitative mating assays were performed as previously described (Peterson et al). In brief, *MATa* or *MAT alpha* strains (*ahis4* and *alpha his4*) were diluted to 2 OD/ml and 400ul of the strains were plated onto YMD plates. 10-fold serial dilutions of tester strains (1.0 to 0.01 OD/ml) were spotted on mating lawns. Mating is only able to occur when the *MATa* genes at *HMR* are repressed. Mating leads to complementation of auxotrophic markers in a diploid and growth on YMD plates (lacking any supplements).

### Quantitative Mating

Tester strains were grown in YPD for 12-14h at 23° C. The mating lawn strain (*MATa his4*) was grown in YPD at 23° C for 12-14h. Cell density was measured at OD600. The mating lawn strain was centrifuged and resuspended in fresh YPD to a concentration of 2 OD600/ml. Tester strains were diluted to 1 OD600/ml and then serially diluted 1:10 to an absorbance of 0.0001OD/ml. 100ul of each dilution was mixed with 300ul of mating lawn and all 400ul were plated onto YMD plates lacking supplements. In parallel, 300ul diluted tester strains were mixed with 100ul YMD and plated onto YMD plates containing all supplements (as controls). Cells were allowed to grow on the plates at 30° C for 3 days and colonies were counted. Three independent colonies of each strain were analysed.

### Fluorescence Imaging

Microscopy was performed on live cells as described ^70^. Cells were grown exponentially in YMD (yeast minimal dextrose) + amino acids (Leu, Ura, Trp, Lys, Ade, and His) to an OD600 nm of ∼0.6. Cells were rinsed in YMD + amino acids and placed on YMD + amino acids and 1.5% agarose patches on slides, covered with a microslip, and imaged. Images were acquired on an inverted wide-field microscope (Xi70; Olympus) with precise stage (DeltaVision; Applied Precision) using a camera (CoolSNAP HQ2; Photometrics). Optical image stacks of 20 images were acquired with a step size of 200 nm for 400–500 ms in the appropriate wavelength channel (CFP/YFP/GFP/mCherry). A 100Å∼/1.4 NA oil objective was used. Acquisition software softWoRx 3.7.1 (Applied Precision) was used for image acquisition and analysis. Cropping of images was performed in Photoshop (Adobe).

Multiple independent trials were performed for each strain and approximately 100 cells were analysed in each trial for each strain.

### Fluorescence Cytometry

Yeast cells were transferred from a plate using a frogging tool (Sigma) into three different microtiter plates each containing 100ul YMD+ all supplement media. Cells were grown overnight at 30° C without shaking.

10 microliters of each culture were used to inoculate deep well (2 ml) microtiter plates containing 590 ul YMD+ supplements (Leu, Ura, Trp, Lys, Ade, and His). Cultures were grown overnight at 30° C with shaking at 600 rpm. 10 microliters of these overnight cultures were used to inoculate fresh 2ml microtiter plates containing 590 ul YMD+ supplement media and grown overnight at 30° C with shaking. The overnight cultures were diluted with fresh YMD+ supplement media (500ul) in 2ml microtiter plates to an OD/ml of 0.2 and grown at 30° C for 3-4 h with shaking. 400 microliters of each culture were removed and filtered through a nitex screen and fluorescence was measured.

For induction experiments, cells were grown in YMD media containing 3% dextrose with supplements and either with or without methionine.

The ATTUNE NxT flow cytometer with lasers BRV6Y was controlled by the ATTUNE cytometric software version 5.3.0. Approximately 10,000 cells were recorded per sample. Analysis of the data was performed using Flow Jo. Initial gating was of forward scatter as well as side scatter representing the distribution of cells based on their intracellular composition and size. This was followed by the drawing of a Boolean AND gate prior to the measurement of fluorescence. Each biological replicate was analysed separately multiple times.

### Multi-Focus Microscopy

Cells were grown exponentially in yeast minimal medium with appropriate supplements and 2% dextrose (YMD+all) on a roller drum at 30°C overnight. The cultures were then back-diluted to an OD600 of 0.2/mL in YMD+all and returned to the roller drum at 30°C for 4 hours. Cells were then pelleted from 1 mL of culture and resuspended with 20 µL of YMD+all. 3 ul of this suspension was applied to a 1.5% agarose YMD+all pad on top of a microscope depression slide and cover slipped.

Images were taken on a high-resolution multifocus microscope as described previously ^71^, which allowed for the simultaneous acquisition of two-dimensional images across 9 different focal planes spaced 144 nm apart. The diffraction limited axial resolution of the ∼525nm fluorophore was around 600nm with a 1.4NA. The lens is a silicon immersion Olympus 1.3NA 60x lens and the total magnification is around 180x, diffraction limited to ∼200nm. Using a two-z-step acquisition for each time point, we were able to capture 18 focal planes of cells across a depth of 3 um per image with minimal photobleaching. For each field of view, an image was captured every 15 seconds over a 10-minute period, totaling 41 time points.

Image analysis was performed using the FIJI distribution of ImageJ software ^72^. The 18 images across 41 time points were first re-stacked with a MATLab script and ImageJ. Processing of the images initially involved bleach correction by the ImageJ plugin using the histogram matching method ^73^. To enhance the contrast between signal and background, maximum intensity and standard deviation Z projections of these bleach corrected images were then prepared and multiplied against each other. Spots and tracks were identified, a threshold was applied and curated against false positives via the TrackMate plugin ^74,75^. To obtain the original maximum intensity measurements before bleach correction, a copy of the TrackMate overlay data were saved and edited such that TrackMate redirected and performed the annotations on a maximum intensity Z projection of the re-stacked file without any other processing.

## Results

### Construction of yeast reporter strains to measure silencing

To systematically measure the potential susceptibility of various enhancers and promoters to heterochromatin silencing, we expanded the modular system developed recently to study gene activation ^76,77^. This allowed us to generate various permutations of regulatory elements to investigate the functional relationship between silencers, UAS enhancers and core promoters (Figure 1A). The system we generated involves silencers flanking coding regions (CDS) of various reporters – *MATa1*, *URA3*, Venus+ PEST+NLS, mCherry+ PEST+NLS and 14xPP7-*PHO5*. Immediately upstream of the CDS, we inserted UAS enhancer + core promoter segments (*MATa1, PDC1, PGK1, RPL28, TDH3, ACO1, CDC19, ADE2*, *GAL1 and MATalpha2)* that we had previously characterized for gene activation ^77^. In this manuscript the term UAS enhancer is defined as a genetic element that interacts with specific activator proteins to stimulate transcription from a separable core promoter. We define a core promoter by its occupancy by GTFs based on ChIP-seq data ^78^. While the transcription rate mainly determines the mRNA abundance in yeast, modulation of mRNA stability can also be a factor ^79^ and the 3’UTRs play a role in mRNA stability and mRNA abundance ^77^. To minimize this variable in our analysis we placed the *PGK1* 3’UTR and transcription terminator downstream of all the reporter CDS. These constructs were integrated at one of two sites in the yeast genome using Cas9+CRISPR; the *HMR* locus near the right telomere of chromosome III or at the *LEU2* gene which is approximately in the middle of the left arm of chromosome III.

**Figure 1:**
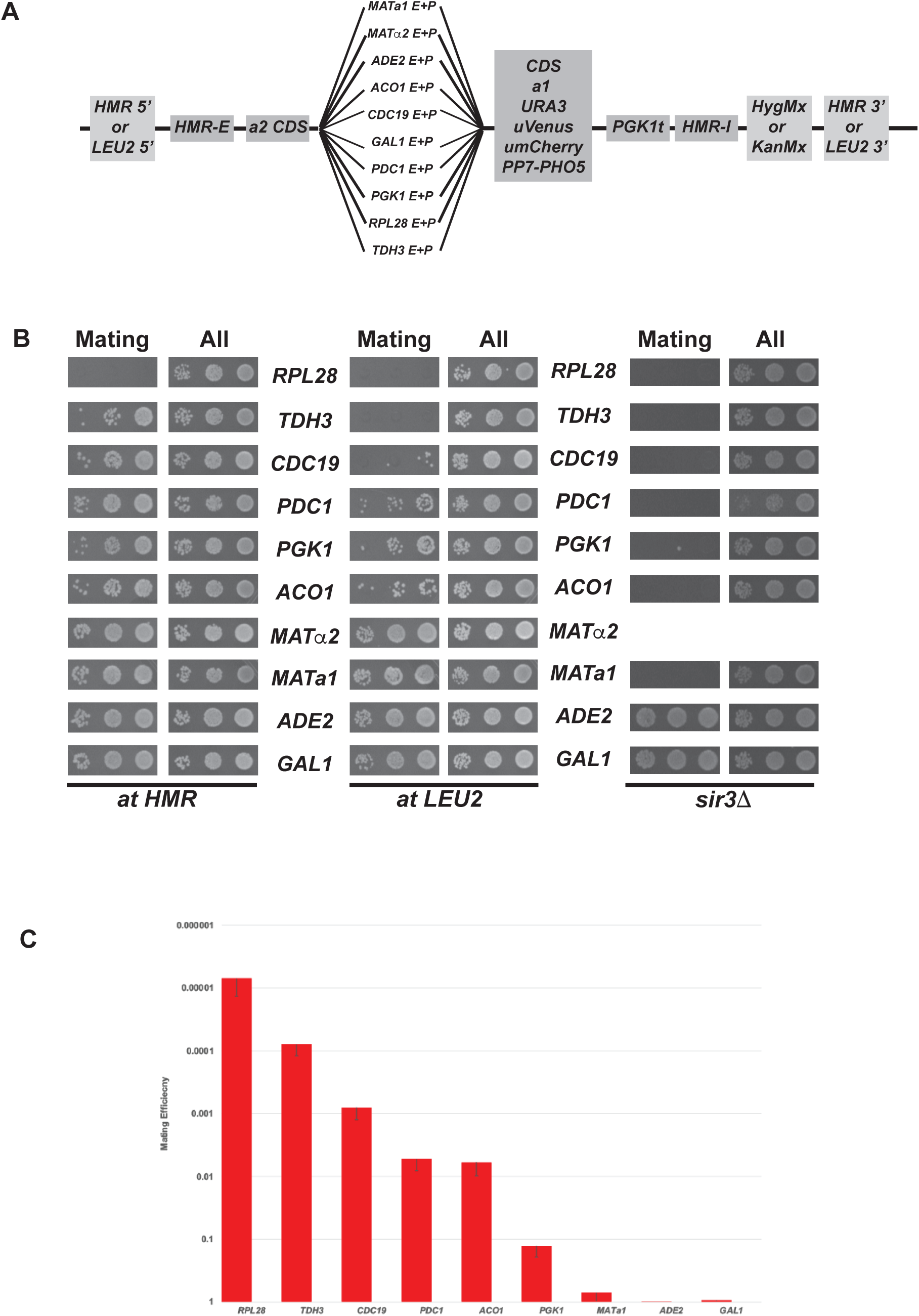
Measuring silencing with the *MATa1* CDS. A) Schematic of strain constructions. Schematic representation of the silencing cassettes constructed from permutations of different regulatory elements and reporter genes. All cassettes were integrated at either the *LEU2* or *HMR* loci on chromosome III. B) Left panels: Repression of the wild type *HMR* cassette located at *HMR* monitored by mating assays. Strains contain the *MATa1* reporter under the control of different regulatory sequences. Ten-fold serial dilutions of the various strains were spotted on YMD+all mix or plates with appropriate mating type tester lawns. Colonies were allowed to grow on plates for 2-3 days prior to documentation. Middle panels: Repression of the wild type *HMR* cassette located at *LEU2* monitored by mating assays. Strains contain the *MATa1* reporter under the control of different regulatory sequences. Cells were grown and diluted as described above. Right panels: Repression of the wild type *HMR* cassette monitored by mating assays in *sir3* delete strains. Strains contain the *MATa1* reporter under the control of different regulatory sequences. Cells were grown and diluted as described above. C) *Q*uantitative mating of the wild type *HMR* cassette located at *LEU2.* Quantitative mating analyses of various regulatory elements driving expression of *MATa1* were performed as described previously ^126–128^. The data are presented as diploid CFU and are mean values from three independent experiments carried out in parallel.

### HMR only partially represses housekeeping gene enhancers/promoters

A haploid yeast strain of the alpha mating type will mate with cells of the opposite mating type to form diploid cells only if the *MATa1* gene at the silenced locus is transcriptionally repressed. Derepression of the *MATa1* gene renders haploid alpha mating type cells unable to mate and form diploid colonies. We built constructs where the *HMR* locus with its native *HMR-E* and *HMR-I* silencers flanked the *MATa1* CDS fused to the ten different enhancers and promoter sequences, and these were integrated at the *HMR* locus. The strains were analysed using a semi-quantitative mating assay (Figure 1B). The data show that the *MATa1, MATalpha2, ADE2* and *GAL1* enhancer/promoters were efficiently repressed. Amongst the other regulatory elements, there was partial but varying degrees of repression of the *ACO1, PGK1, PDC1, CDC19* and *TDH3* elements while the *RPL28* regulatory elements resisted silencing almost completely.

The enhancer and promoter elements that were partially derepressed at *HMR*, became further derepressed when these silencing cassettes were instead integrated at the *LEU2* locus. *ACO1, PGK1* and *PDC1* regulatory elements were derepressed to a greater extent and the *TDH3* and *CDC19* regulatory elements resisted silencing completely (Figure 1B). The four cassettes that were fully repressed at *HMR* remained repressed at *LEU2* and these results indicate that robust silencing is position dependent.

Quantitative mating analysis revealed a gradient of repression dependent on the enhancer and promoter sequences regulating *MATa1* CDS at *HMR.* The *MATa1, ADE2* and *GAL1* regulatory elements were silenced in synthetic complete media in most cells, whereas the *PGK1* enhancer/promoter was repressed in ∼10% of the cells, *PDC1* and *ACO1* were repressed in ∼1% of the cells, *CDC19* showed repression in only 0.1% of the cells while the numbers were even lower for *TDH3* and *RPL28* enhancers/promoters (Figure 1C).

The complete or partial repression observed with these constructs was Sir dependent. In a strain lacking Sir3, repression was completely lost for all the enhancers and promoters tested except for *ADE2* and *GAL1* (Figure 1B). This is unsurprising, because silencing was monitored in synthetic complete media where both *ADE2* and *GAL1* genes are known to be transcriptionally inactive even in the absence of Sir proteins.

### Cytometry shows that partial repression of housekeeping gene enhancers and promoters is not bimodal

Based on classic studies on the *MAT*, *URA3* and *ADE2* enhancers/promoters, silencing is believed to be an all or nothing phenomenon exhibiting bistable expression states ^67,80^. The partial repression observed for *TDH3, PGK1, PDC1, ACO1, CDC19* raised the question of whether cells with these elements also exhibited bistable expression states. Cytometry is well suited to analyse variation in gene expression in a cell population. To investigate the partial silencing, we built strains where the Venus CDS containing a nuclear localization signal (NLS) and a PEST sequence was linked to the different enhancers/promoters and we integrated the cassette at the *HMR* locus. We chose to use the fluorescent protein reporter Venus to analyse gene repression because it has a bright fluorescence and matures rapidly ^81^. The presence of a *CLN2* PEST sequence results in a high turnover rate (∼25 min) while the presence of a NLS leads to the concentration of the fluorescent protein in the nucleus ^82–84^. Cells were grown in complete synthetic media and logarithmically growing cells were analysed for expression of Venus after appropriate gating using a flow cytometer. We demarcated cells as being silent relative to expression values of the *MATa1* enhancer/promoter where greater than 99% of these cells were repressed (Figure 3A). For the other enhancer/promoter cassettes, cells with fluorescence values within the *MATa1* regulatory element repressed gate were considered repressed while the remainder of the cells were considered not silent and based on this gating parameter, we quantified the population frequency of cells that were not silenced. The frequency of non-silenced cells was plotted for three independent colonies (Figure 2 top panel). The data show that the *MATa1, ADE2* and *GAL1* enhancers/promoters were fully repressed while partial repression was seen for the other enhancers/promoters with values ranging from ∼20% active to ∼90% active.

**Figure 2:**
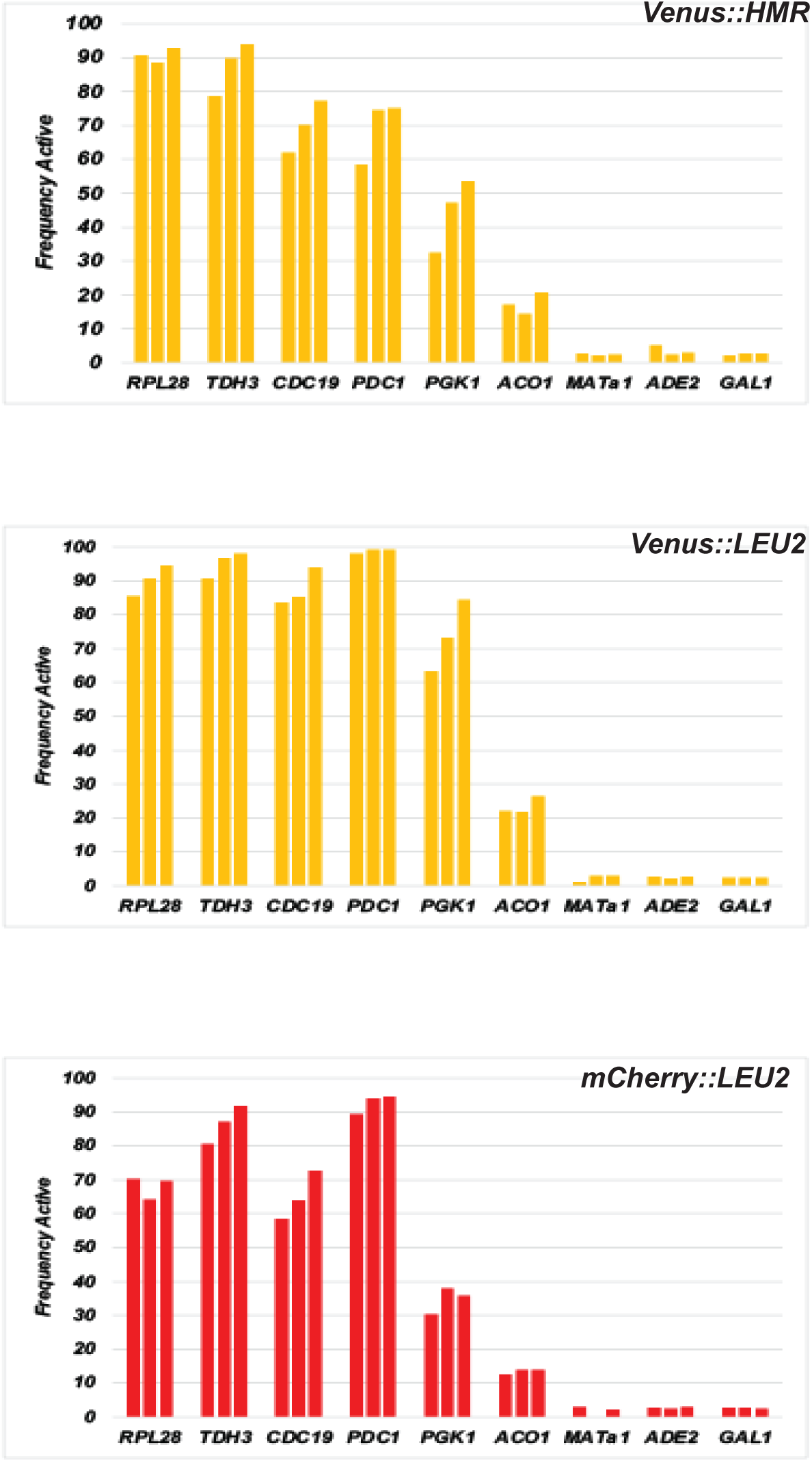
Measuring silencing using different fluorescent reporter proteins. Top panel: Repression of the wild type *HMR* cassette located at *HMR* monitored by fluorescent cytometry using the Venus reporter gene. Different UAS enhancers and core promoters were linked to the Venus reporter gene and the silenced cassette was integrated at the *HMR* locus. Expression of the constructs was measured using a flow cytometer. The frequency of cells that escaped silencing were determined using the *MATa1* regulatory element as a standard. Strains with the *MATa1* enhancer and core promoter were used to set the silenced gate such that 99% of cells were silent for this cassette (see figure 3). Three separate colonies were measured for each strain and the data from each were plotted separately. Middle panel: Different UAS enhancers and core promoters were linked to the Venus reporter gene and the wild type *HMR* silencers were integrated at the *LEU2* locus. Expression of the constructs was measured using a flow cytometer as described above. Bottom panel: Different UAS enhancers and core promoters were linked to the mCherry reporter gene and the wild type *HMR* silencers were integrated at the *LEU2* locus. Expression of the constructs was measured using a flow cytometer as described above.

**Figure3:**
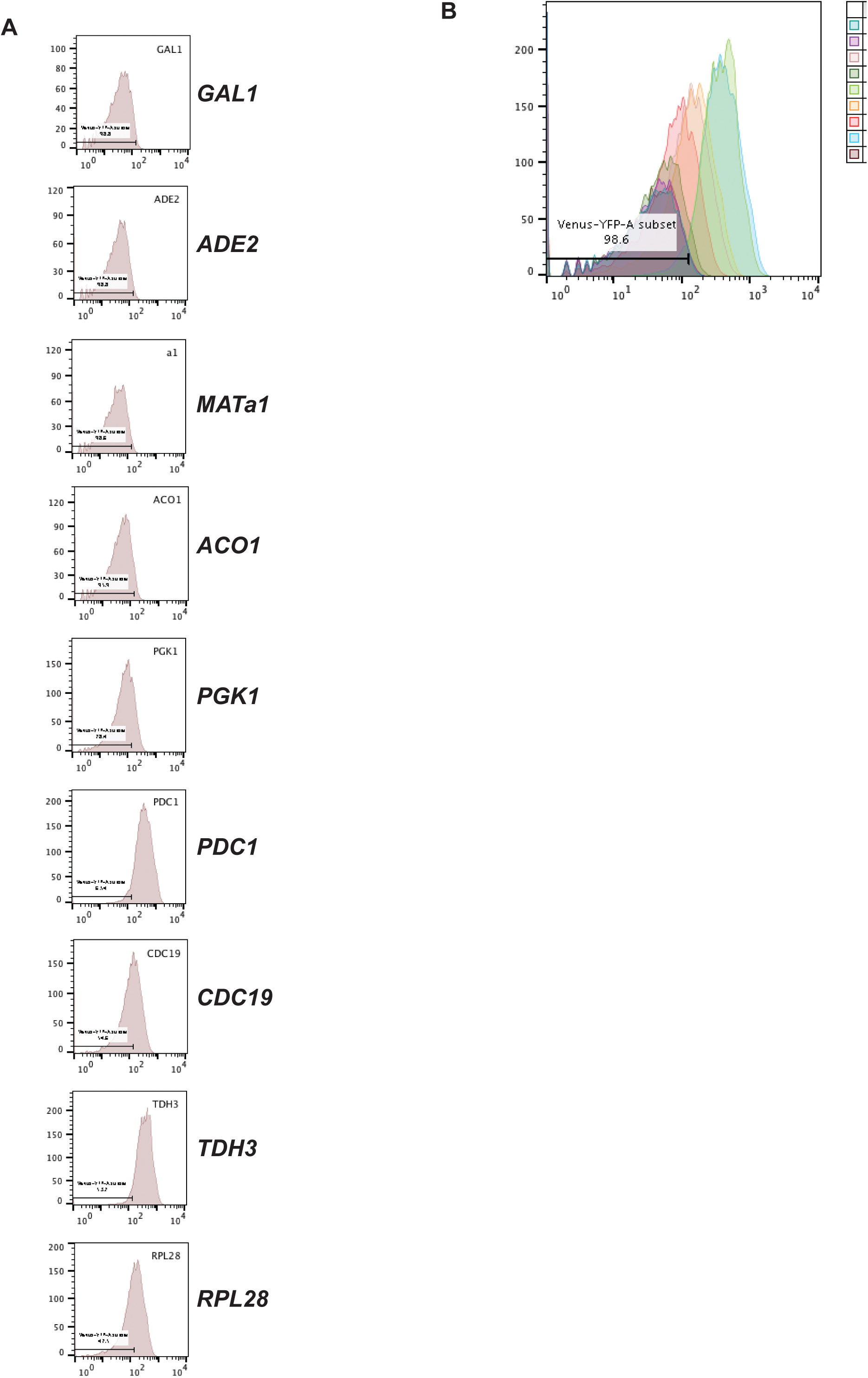
Histogram of fluorescent cytometry with wild type *HMR* and the Venus reporter. A) Histograms of the fluorescence cytometry analysis of the expression of Venus under the control of different enhancers and core promoters at *HMR* were generated. X-axis corresponds to fluorescence levels and Y-axis to the total cell counts. B) Overlay of the different histograms of the fluorescent cytometry data showing a continuum of expression values.

When the same analysis was done with *HMR* cassettes containing Venus integrated at the *LEU2* locus we observed similar results but with reduced frequency of repression of the different cassettes (Figure 2 middle panel). These results are consistent with the mating assay showing position dependent repression levels. Similar results were observed when the Venus reporter was replaced with the mCherry reporter (Figure 2 bottom panel) and when the *CLN2* PEST sequence was eliminated (data not shown) suggesting that this pattern is not primarily a function of protein turnover rates.

The protein expression profiles of individual cells in a population can also be informative with regards to mechanisms of regulation. Monitoring the expression histograms of these cells undergoing silencing did not show a bimodal peak profile that would be expected for bistable expression states (Figure 3A). The histograms show a single peak of expression though for the house keeping genes we observe a skewed normal distribution with a lagging edge extending towards the repressed state. Superimposing these peaks showed a continuum of protein expression values for the different enhancers/promoters indicating that the partial repression observed was not an all or nothing phenomenon, but the amount of fluorescent protein within individual cells were present at varying levels (Figure 3C). We are therefore unable to distinguish transient repression from stable silencing based solely on these cytometry profiles.

### Partial repression of housekeeping gene enhancers/promoters is not stably inherited

The *URA3* gene is commonly used as a reporter to measure Sir-mediated silencing ^80^. Repression of *URA3* measured by colony formation on medium lacking uracil or containing 5-FOA measures repression throughout the cell cycle and over multiple generations. Stably inherited repression of *URA3* allows cells to form colonies on medium containing 5-FOA and is a signature of an epigenetic silenced state. We therefore built the *HMR* cassette with the different enhancer/promoters driving expression of the *URA3* CDS. These cassettes, integrated at *LEU2*, were monitored by growth on media lacking uracil or media containing 5-FOA. All strains grew normally in synthetic complete media (Figure 4). In media lacking uracil, we did not observe any growth for strains with the *GAL1* enhancer/promoter, very weak growth for the *ADE2* enhancer/promoter and robust growth for the *MATa1, MATalpha2, ACO1, PGK1, PDC1, CDC19, TDH3* and *RPL28* enhancer/promoter. On the other hand, in media containing 5-FOA, we observed robust growth for strains with the *GAL1, ADE2*, *MATa1* and *MATalpha2* enhancer/promoter but no growth for the remaining six-*ACO1, PGK1, PDC1, CDC19, TDH3* and *RPL28* enhancer/promoter. These results indicate that the partial repression observed for the housekeeping gene (*ACO1, PGK1, PDC1, CDC19, TDH3* and *RPL28)* enhancer/promoter is not stably inherited at all. On the other hand, *MATa1, MATalpha2* and *ADE2* growth on both - uracil and 5FOA media is typical of bistable expression states as has been previously reported. These data indicate that stability of silencing is not simply a function of silencer and silencing proteins but also influenced by the strength and/or property of the enhancer/promoter element undergoing repression and possibly of the gene product being used as the reporter in the silencing assays.

**Figure 4:**
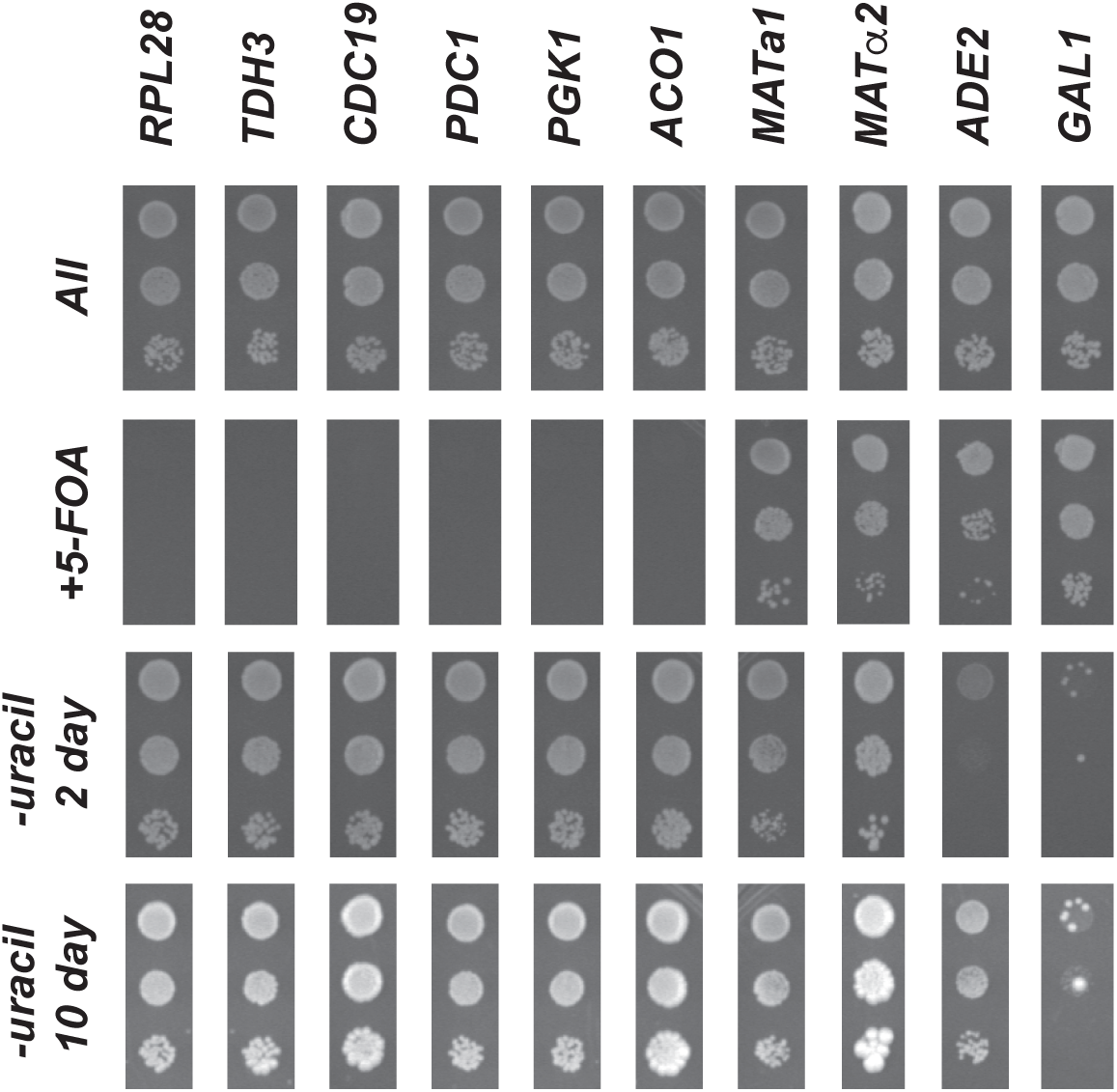
Measuring silencing of the wild type *HMR* cassette with the *URA3* CDS. The *URA3* reporter was linked to various regulatory elements and the *HMR* cassette was integrated at *LEU2*. Expression of the gene was monitored by growth on YMD plates lacking uracil or containing 5-FOA. Approximately 3 ul of ten-fold serial dilutions of overnight cultures was spotted on the different plates. Cells were allowed to grow for 2 to 10 days before the plates were photographed.

### Measurement of Transcription Foci in cells undergoing Silencing

Sir mediated gene silencing operates at the level of transcription but the different assays we have used so far (*MATa1, URA3* and *Venus*) analyse protein/enzyme function, which is a step removed from transcription. We therefore decided to directly assay mRNA synthesis in live cells and visualize transcription initiation. We utilized an array of 14 binding sites for the bacteriophage coat glycoprotein PP7 and inserted these repeats in the 5’UTR of the *PHO5* CDS ^43,49^. This construct was linked to the different UAS enhancers and core promoters in the silencing cassette. The transcribed mRNA forms multiple stem loops in the 5’UTR, allowing the bacteriophage protein PP7 to bind. The nascent mRNAs can thus be tracked in individual cells that are constitutively expressing the PP7-GFP fusion protein in real time using a sensitive high-resolution wide-field fluorescence microscope (Figure 5). With this construct we can directly visualize and quantify loci that are not silent and are instead undergoing mRNA synthesis in live cells ^49^. We quantified the number of PP7-GFP transcription foci at *HMR* in approximately 400 cells grown in synthetic complete media and found that the number of transcription foci correlated with the Venus expression profiles we had observed in our flow cytometry assay. When the same analysis was performed in a Sir3 delete strain, the number of expression foci increased two to three fold for the moderately repressed *PDC1, PGK1* and *ACO1* cassettes and increased to a lesser degree or not at all for the very strong regulatory elements (*TDH3, RPL28* and *CDC19*) suggesting that these regulatory elements were undergoing robust transcription even in the presence of Sir proteins. We did not observe any transcription foci in wild type cells for the *MATa1*, *ADE2* and *GAL1* regulatory elements.

**Figure 5:**
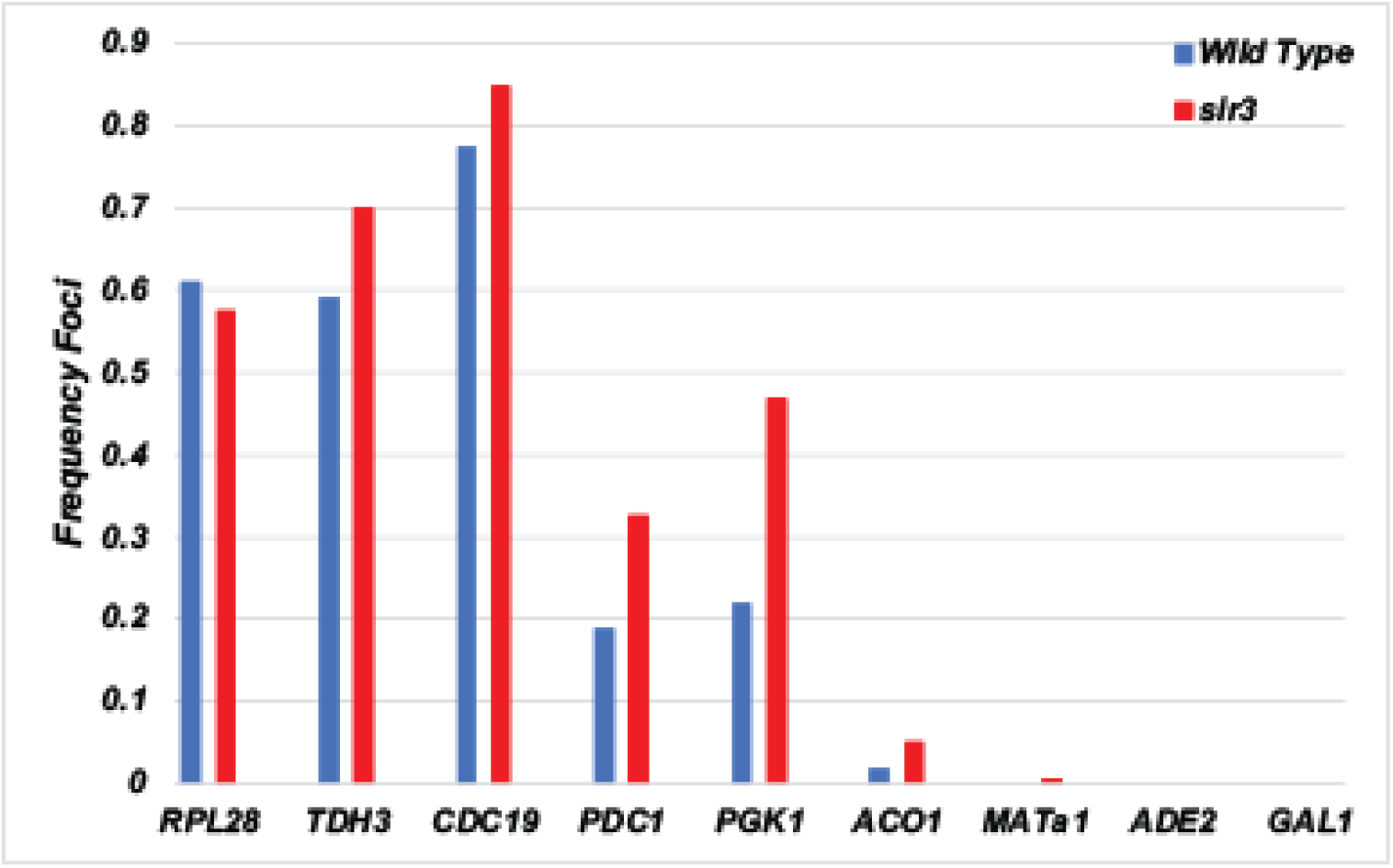
Real-time measurement of fluorescent mRNA synthesis using wide-field fluorescent microscopy. A) The reporter gene (*PP7bs-PHO5*) was linked to various regulatory elements and the silenced cassette was integrated at *HMR* in a strain constitutively expressing PP7-GFP. mRNA synthesis (GFP foci) was monitored using a fluorescence microscope. All images are the sum of the different projected z-stacks. Approximately 400 cells were analysed, and each strain was analysed over multiple days. The number of fluorescent foci were quantified as a percent of the total number of cells analysed and plotted for wild type cells as well as cells lacking Sir3.

### Measuring Duration of Transcription Bursts

Measuring transcription foci quantifies the number of cells where the *HMR* locus is derepressed but is not very informative with regards to the duration of time when the locus is active. To measure duration when the *HMR* locus in active we resorted to the use of a multi-focus microscope (MFM) and coupled this instrument with the *PHO5*-PP7-GFP system. This system allows us to visualize transcription initiation and elongation in live cells and determine the frequency and duration of transcription of a gene over a period of time. The MFM simultaneously acquires images of multiple focal planes at a single moment in time thus reducing photobleaching and allows the repeated imaging of a single cell over an extended period of time. However, previous work using this system has shown that the lower exposure time and reduced intensity of the laser only allows the detection of transcription foci with three or more engaged RNA polymerases ^49,85^. Therefore, the presence of fluorescent foci using the MFM only identifies loci with multiple transcriptionally engaged RNA polymerases, and the numbers obtained are an underestimate of the actual number of RNA polymerases transcribing the locus in any given cell. While not optimal, this system allows us to differentiate periods when the locus is highly active from periods when the locus is less active or inactive (Figure 6A and 6B). A field of cells were visualized every 15 seconds for 10 minutes and the presence of a transcription focus was highlighted. For the three regulatory elements analysed (*TDH3, PGK1* and *RPL28*) the MFM data identified some cells in the population where we did not detect any transcription over the 10-minute period under observation while there were other cells where we did observe transcription. In cells with a transcription focus, transcription was intermittent suggesting bursting. For each cell, the duration of time that the locus was undergoing transcription was quantified as the frequency of activity and the data were plotted as a boxplot. The same analysis was also performed on *sir3*Δ strains and these data show an increase in transcription in cells lacking Sir3 compared to wildtype cells (Figure 6C). The loss of Sir3 led to an increase in the total time when fluorescent foci were observed in all three strains though the effect was less pronounced for the *RPL28* regulatory sequences. Since the presence of a detectable fluorescent focus indicates a cell with multiple engaged RNA polymerases these data collectively suggest that Sir3 likely functions by reducing the time when the promoter is open and accessible and suggests that this protein likely functioned by blocking the repeated and frequent initiation of transcription by RNA polymerase thus reducing burst durations.

**Figure 6:**
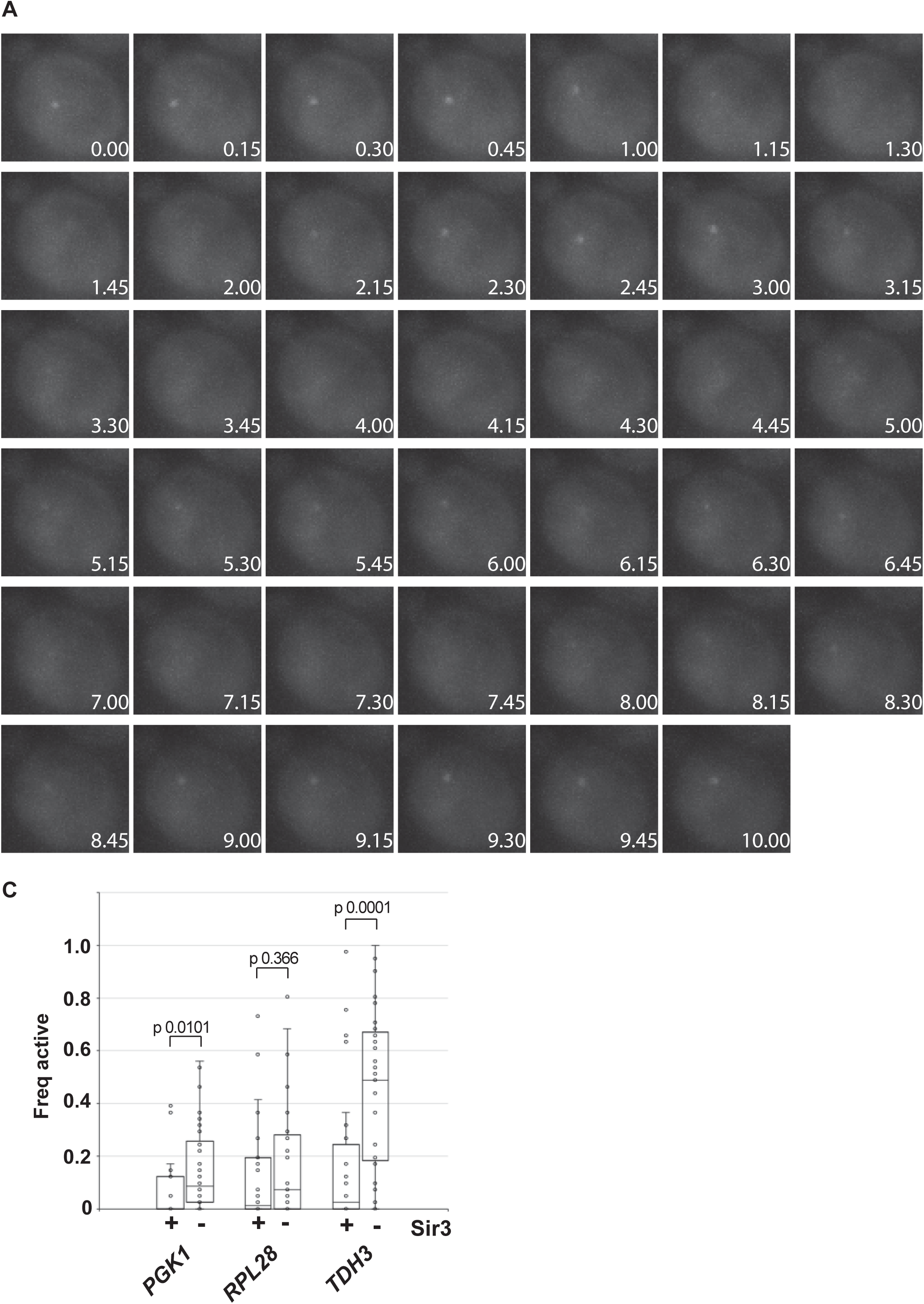
Real-time measurement of mRNA synthesis using a multi-focus fluorescent microscope. A) Time lapse images of a yeast cell undergoing silencing. A single yeast cell was imaged every 15 seconds across 18 different focal planes using a multi-focus fluorescent microscope. The appearance or disappearance of the focal spot was monitored over a 10 minute time period. B) A GIF file of the 41 images. C) Length of time when a locus was transcriptionally active in each cell. The duration of time that the *HMR* cassette was transcriptionally active in individual cells was determined from fluorescence dot intensity traces and plotted as a box plot. The fluorescent foci were monitored every 15 seconds in individual wild type and *sir3*Δ cells over a period of 10 minutes. The fluorescence intensity of the dots was plotted as a function of time. Values above background were considered transcriptionally active.

### Sir mediated repression operates on both enhancers and promoters

Some studies have indicated that the Sir proteins block access to sequence specific transcription activators that bind UAS enhancers while other studies suggest that the Sir proteins block general transcription factors from binding the core promoter sequences ^60,63,86,87^. To measure the individual contributions of the UAS enhancer and the core promoter in resisting repression, we adopted the approach we used previously to delineate and characterize the strengths of the enhancers and promoters in gene activation ^77^. We built a matrix of seven enhancers linked to seven core promoters and asked if the extent of repression of these 49 constructs was solely a function of the enhancer sequences, solely promoter dependent or depended on both elements. Fluorescence cytometry measurements of the Venus reporter showed that both the enhancer and the core promoter influenced expression to different degrees (Figure 7A). Enhancer strength clearly was a major determinant in the ability of that regulatory element to resist Sir mediated repression. The *TDH3* enhancer was unable to be silenced to a large extent when linked to different promoters as were the *RPL28* and *CDC19* enhancers while the *PGK1* and *ACO1* enhancers had intermediate ability. However, what was also clear from these analyses was that certain core promoters significantly influenced the ability of these strong enhancers to resist repression. For example, repression of the *TDH3* enhancer was significantly increased when this element was linked to the *MATa1* or *ACO1* core promoters indicating that sequences within these two core promoters were particularly susceptible to being transcriptionally repressed suggesting that underlying promoter sequences play an important role in regulating expression levels of loci undergoing Sir mediated repression.

**Figure 7:**
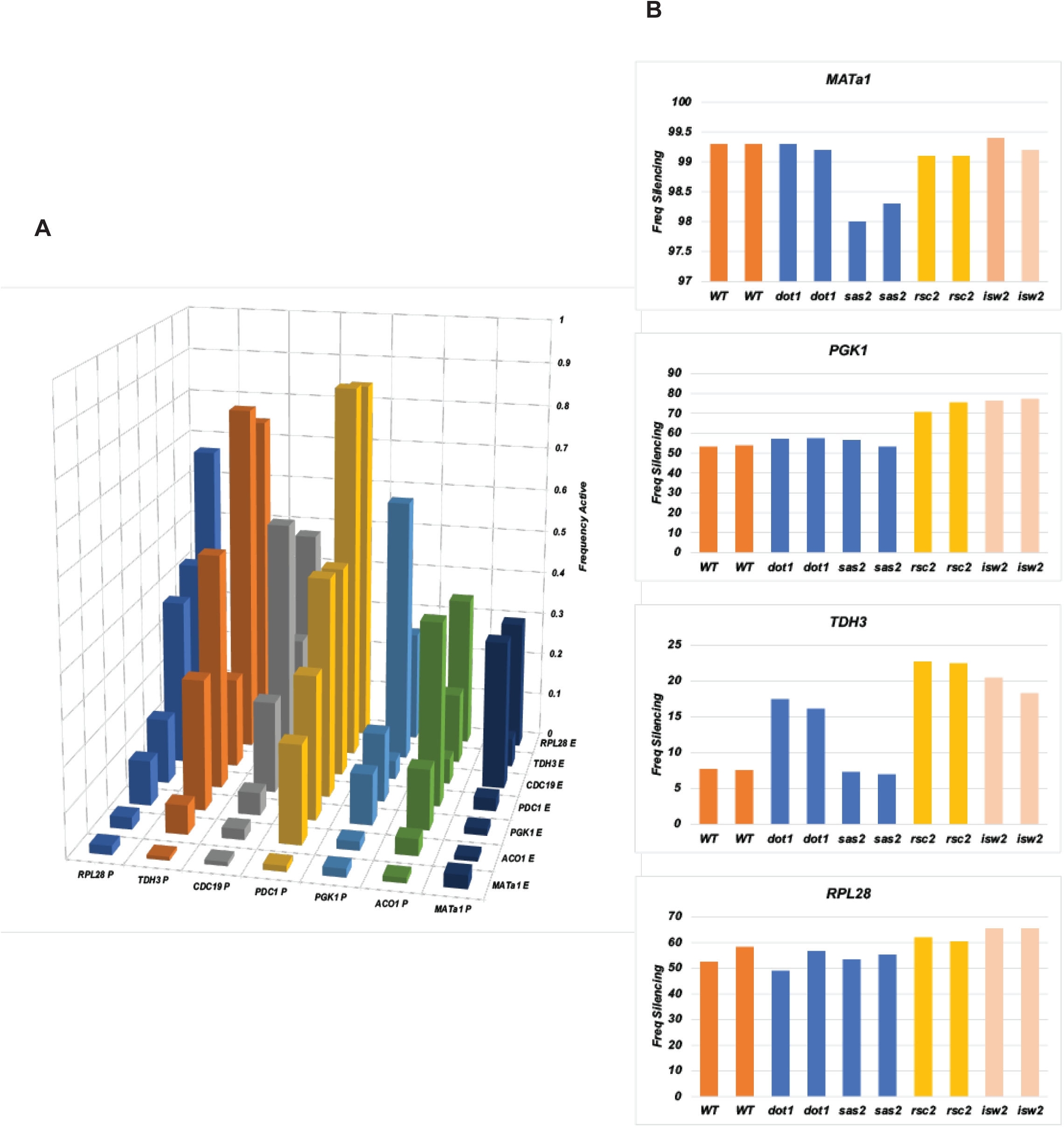
Expression analysis of regulatory elements controlling Venus’s expression at *HMR*. A) A 7x 7 matrix of different combinations of UAS enhancers and core promoters with the Venus reporter gene and the *PGK1* 3′ UTR was generated. These constructs were flanked by *HMR* wild type silencers. Venus’s expression of these constructs was measured using a flow cytometer. Three separate colonies were measured for each construct and the average values were plotted. B) Fluorescent cytometry measurements of wild type *HMR* with a Venus reporter in strains lacking Dot1, Sas2, Isw2 and Rsc2. Expression of *HMR* loci containing different UAS enhancers and core promoters linked to Venus was measured by fluorescence cytometry. Bar graphs of Venus expression of wild-type and various mutant cells-*sas2Δ, dot1Δ, isw2Δ and rsc2Δ* grown in YMD medium are plotted for four different UAS enhancers and core promoters. Each measurement was performed with two colonies. Please note that the Y-axis scales are different in each of the graphs.

### Loss of Specific Chromatin Remodellers Increases Gene Repression

One mechanism by which regulatory elements affect transcription is via their ability to exclude or favour nucleosome formation since most transcription factors cannot bind their recognition sites if those elements are packaged into nucleosomes ^22,23^. Nucleosome presence at regulatory elements is affected by histone modifiers and chromatin remodellers. In wild type cells the actions of these enzyme complexes are to create a chromatin architecture at the core promoter that is nucleosome depleted thus favouring transcription. We built strains where subunits of either chromatin remodellers or histone modifiers *RSC, ISW2, SAS-I* and *DOT1* were deleted. We monitored expression of four different regulatory elements (*MATa1, PGK1, RPL28, TDH3*) driving expression of *HMR*::Venus (Figure 7B). Our data show an enhancer/core promoter dependent effect. The *RPL28* regulatory sequences remained resistant to gene silencing even upon loss of various cofactors. This is consistent with previous data showing that the NDR at ribosomal protein coding genes is mediated primarily by the GRFs ^88^. In contrast, *TDH3* and *PGK1* regulatory sequences are dependent upon both RSC and ISW complexes in resisting silencing. Loss of these chromatin regulatory cofactors resulted in an increase in these enhancers and promoters becoming repressed. These results suggest that the formation/maintenance of a specific chromatin architecture was important for efficient repression and was consistent with results showing that alteration in the number and optimal spacing of nucleosomes affects the stability of silencing ^89^.

### Increasing silencer strength enabled repression of strong enhancers and core promoters

Our data with the native *HMR* silencers suggest that the strong housekeeping gene enhancers and promoters fully or partially resist being repressed. One question was whether this was an intrinsic property of these gene regulatory sequences or simply a function of relative balance between activators and repressors. Previous data showed that Sir1 binds *HMR-E* in a sharp narrow peak reflecting localized binding of this protein ^59^. Recent analysis of silencing in Sir1 mutant cells showed that loss of Sir1 led to decreased recruitment of the other Sir proteins to the silenced locus coupled with a small reduction in the spread of these proteins across the repressed domain ^90^. We therefore decided to investigate the effects of increasing the number of Sir1 binding sites at *HMR* silencers. We constructed synthetic silencers where the wildtype ORC binding site of the *HMR-E* silencer was replaced with four binding sites for Gal4, and the ORC binding site at *HMR-I* was replaced with five binding sites for Gal4. We placed the Venus reporter linked to the various regulatory elements between these two synthetic silencers and integrated this cassette at the *HMR* locus (Figure 8A). Cells containing this cassette also expressed Gal4-Sir1 fusion protein under an inducible *MET17* enhancer/promoter such that Gal4-Sir1 is expressed only in media lacking methionine. Expression and binding of Gal4-Sir1 to the synthetic silencer cassettes led to robust repression of all regulatory elements, though the *RPL28* regulatory element was repressed to a lesser degree compared to the other elements (Figure 8B). These results indicate that even the strongest enhancers and core promoters can be repressed, provided the amount of Sir1 bound to the silencer is increased significantly.

**Figure 8:**
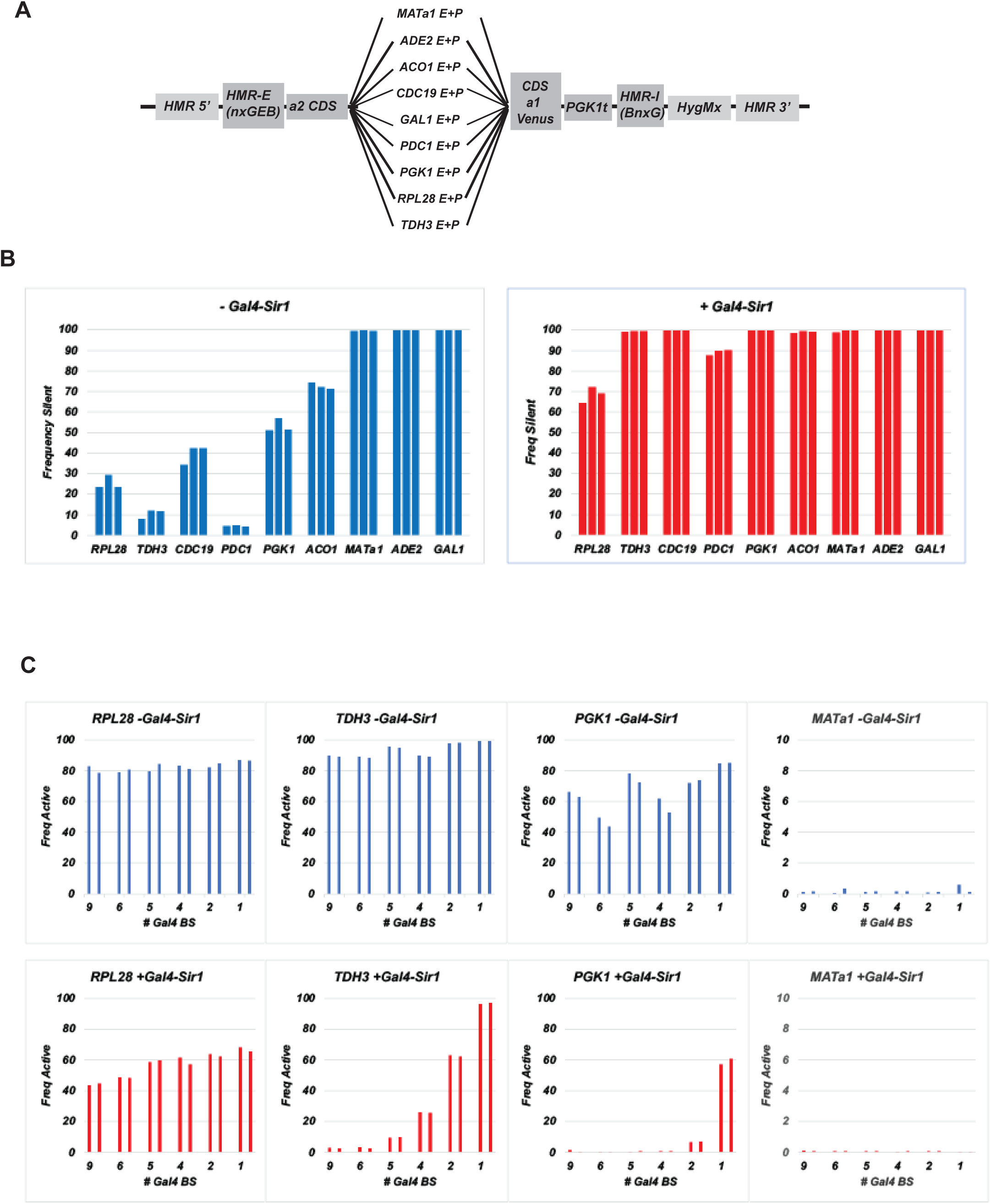
Gal4-Sir1 mediated silencing of synthetic silencer containing *HMR* cassettes. (A) Schematic of the modified silenced locus used in this study resulting in silencing of the Venus reporter gene. (B) Repression of the *HMR* cassette containing four Gal4 binding sites at *HMR-E* and five binding sites at *HMR-I* were monitored by fluorescent cytometry using the Venus reporter gene. Different UAS enhancers and core promoters were linked to the Venus reporter gene and the *PGK1* terminator and the silenced cassette was integrated at the *HMR* locus. Expression of the constructs was measured using a flow cytometer as described. The different strains had Gal4-mRuby-Sir1 under the control of the *MET17* enhancer and promoter integrated at the *his3* locus. Gal4-Sir1 expression was regulated by growing cells in the presence and absence of methionine in the media. (C) Repression of the *HMR* cassette containing variable numbers of Gal4 binding sites at *HMR-E* and *HMR-I* were monitored by fluorescent cytometry using the Venus reporter gene. The *HMR-E* silencer had either four or one binding site for Gal4 while the *HMR-I* silencer had either five, one or zero binding sites for Gal4. Different UAS enhancers and core promoters were linked to the Venus reporter gene and the *PGK1* terminator and the *HMR* cassette was integrated at the *HMR* locus. Expression of the constructs was measured as described above. Figure 10: Schematic representation of the effects of enhancer, promoter and silencer strength on gene silencing.

We were curious about the quantitative relationship between silencer strength and gene repression. While these elements could be repressed by a silenced locus containing nine binding sites for Gal4-Sir1 (four at *HMR-E* and five at *HMR-I*), we decided to vary the number of Gal4-Sir1 binding sites. We built synthetic silencer constructs with either four or one binding sites for Gal4 at *HMR-E* coupled with *HMR-I* silencers with either five, one or zero Gal4-Sir1 binding sites cumulatively leading to nine, six, five, four, two or one binding sites. We analysed the *RPL28, TDH3, PGK1* and *MATa1* regulatory elements driving the expression of Venus. The data show that as the number of binding sites for Gal4-Sir1 increase there is greater repression in the presence of Gal4-Sir1 compared to in its absence (Figure 8C). These results indicate that a key limiting factor in the ability of the native silencers to repress the strong housekeeping gene regulatory elements is the inability of the native silencers to recruit sufficient amounts of Sir1 and possibly the other Sir proteins.

## Discussion

Heterochromatic gene silencing is considered a stable, heritable, and effective form of gene inactivation mediated by a chromatin structure that inhibits expression of most genes regardless of the transcription activator or RNA polymerase involved ^55,56,91,92^.

The *MAT* genes on chromosome III possess a shallow and narrow NDR in their regulatory sequences and are weakly transcribed, generating a few transcripts per cell cycle ^10,12,79^. The wildtype silencers very effectively silence the *MAT* genes at the silenced *HML* and *HMR* loci. These genes are stably silenced by Sir proteins binding to nucleosomes and preventing them from sliding and/or removal thus blocking the formation of a stable transcription complex.

Reducing levels of the Sir proteins via deletions of Sir1, mutations in the binding sites for Rap1 at the silencers or moving the silenced locus to a euchromatic site (*LEU2*) leads to a bistable expression state. Under these conditions, in a fraction of cells, stable transcription complexes can form at the promoters of the genes, allowing these genes to remain active for extended periods of time (Figure 9). The bistable expression state can revert to monostable silencing upon increasing the levels of Sir proteins ^90^. Bistability is also observed in instances where wild type silencers regulate stress responsive and inducible genes such as *URA3, ADE2* or *HSP82*. These genes are activated by single transcription factors and the bistable state is dependent on the activators binding their cognate sites thus helping form stable transcription complexes at the promoter thus resisting repression ^64,65,68,69,80,93^. In contrast, the housekeeping genes have large complex UAS enhancers. Optimal transcription of these genes requires binding of all of the transcription factors while reduced suboptimal expression is observed when some of the binding sites are mutated ^94^. When these genes are placed at a silent locus, rather than observing monostable silencing or bistable expression states, we observe robust but variable levels of transcription that is subtly affected by Sir proteins-an analog rheostat response. This response is directed by the UAS enhancers and core promoters of these genes. The elements escape silencing to varying degrees possibly due to varying levels of occupancy by the transcription activators. These data collectively suggest that Sir mediated repression in wild type cells is optimized to stably silence only weak regulatory elements.

**Figure 9:**
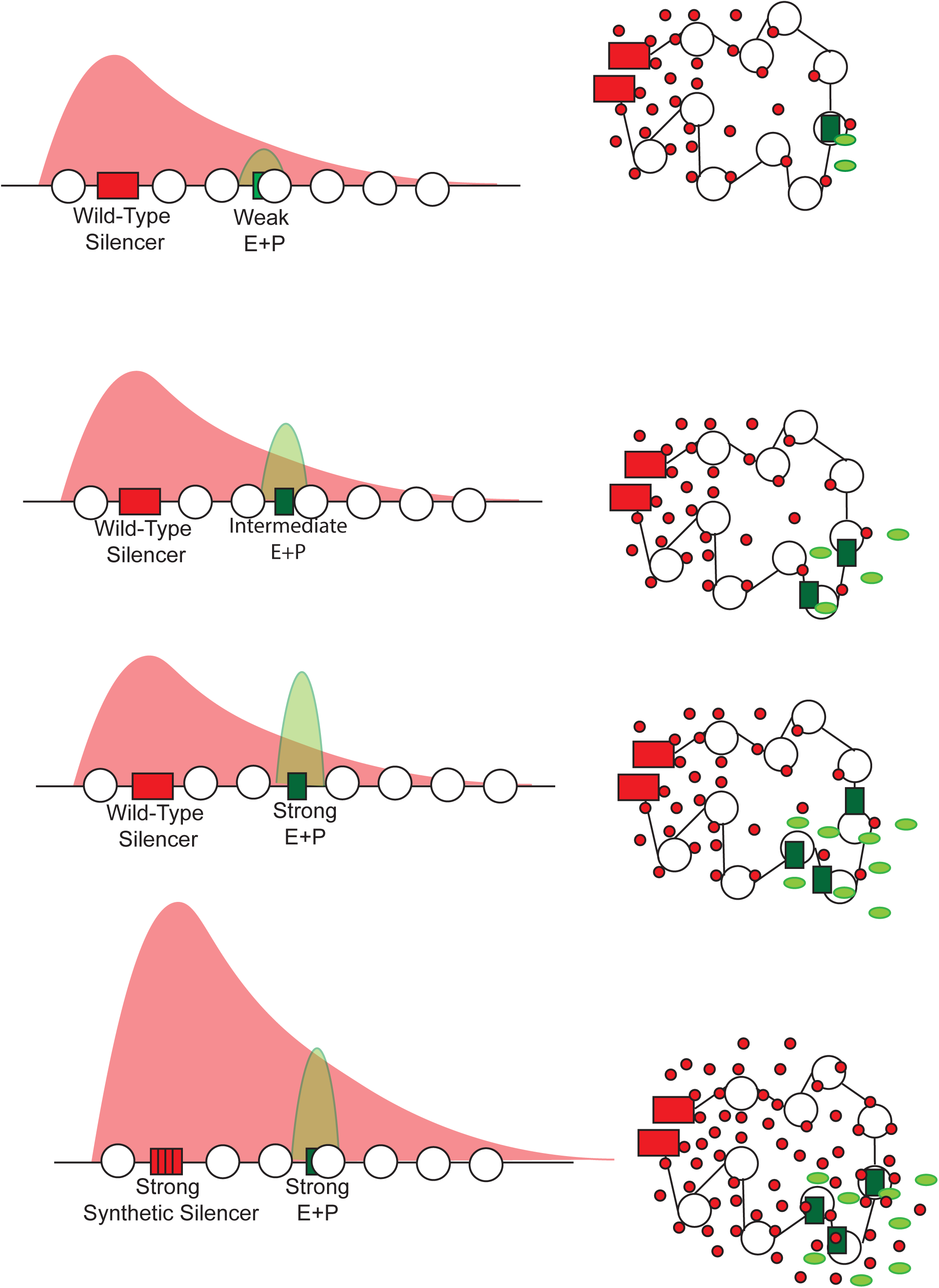
Schematic representation of the effects of enhancer, promoter and silencer strength on gene silencing.

### How might the Sir proteins repress different enhancers and core promoters?

The mechanism by which TFs activate genes is a function of the underlying regulatory sequences as well as the TFs themselves. Different regulatory elements and their attendant TFs have different requirements for histone modifying and remodelling complexes during gene activation. This genetic property of the regulatory system is unlikely to change when a regulatory element is transposed to a silenced domain. It is therefore likely that the Sir proteins negatively influence transcription while being agnostic with respect to theTFs and mechanisms involved. The common step in transcription of most eukaryotic genes is the modification and mobilization of nucleosomes from regulatory sequences. Histone H4 acetylation by modifying enzymes is required for ISW2 ATPase activity and nucleosome spacing by this remodeller ^95^. Similarly, the ability of the Swi/Snf/Rsc remodelling enzymes to engage nucleosomes via their bromodomains is influenced by histone H4 acetylation ^96^. Interestingly, Sir3 interacts with nucleosomes via its Bromo-adjacent homology (BAH) domain making contacts with the unacetylated H4 tail and H2B close to H3K79 ^97^ and these are also the sites required for chromatin remodellers binding. These observations raise the possibility that Sir proteins repress genes by directly affecting the ability of chromatin remodellers to bind to and move nucleosomes thereby altering the temporal kinetics of transcription activator binding, PIC formation and transcription and thus changing the probability mass of active and repressed chromatin configurations. This would be consistent with in vitro studies of RSC and silencing ^98^ as well as with observations showing that Sir bound regulatory regions have significantly lower nucleosome turnover rates compared to euchromatic sites in the genome ^99,100^. This model would also help address a conundrum in silencing which is that while silencing inhibits transcription, the fractal nature of heterochromatin is only subtly different compared to euchromatin and measurements of freely diffusing fluorescent proteins and chemical probes do not show any appreciable decrease in accessibility at heterochromatic loci ^101–103^. This suggests that the nature of heterochromatin mediated inhibition of transcription does not operate at the level of the accessibility of chromatin domains in the nucleus but at the level of the nucleosomal template.

Our analysis of housekeeping genes undergoing silencing demonstrated bursts of transcription (where three or more RNA polymerases are engaged) followed by periods of low activity or quiescence and this was altered in a Sir dependent manner. At the molecular level, we can speculate on how this phenomenon leads to silencing. One could posit that the Sir proteins repress transcription by binding nucleosome and affecting their dynamics. For the very weak and stress induced genes, the presence of nucleosomes over key regulatory sequences and the stabilization of these nucleosomes by the Sir proteins could result in an all or nothing expression phenotype ^87,89^ because the Sir proteins would impede nucleosome mobilization by chromatin remodellers. On the other hand, the housekeeping gene regulatory sequences utilize multiple pioneer transcription factors. Some of these factors recruit chromatin remodellers while others can bind their sites even when these sites are packaged in nucleosomal DNA ^28,88^. The directed induction of active nucleosomal configurations by these TFs allow these regulatory elements to partially or fully resist and even possibly overcome Sir mediated repression.

It is therefore possible that the Sir proteins influence the frequency of chromatin configurations by directly hindering chromatin modifying and remodelling enzymes and indirectly hinder transcription factor binding to regulatory sequences leading to silencing of weak and stress induced genes that are dependent upon these remodelling/modifying complexes.

### How do increased levels of Sir1 at a synthetic silencer induce repression of even the strong housekeeping gene regulatory elements?

The silencers are critical for silencing and loss of these elements immediately abrogates silencing ^104,105^. There are three proteins bound to the silencers and any two of these are sufficient for robust silencing ^106^. The function of Abf1 is to create directionality in silencing via its ability to bind/recruit chromatin remodellers thereby generating evenly spaced nucleosomes ^28,107^. The role of Rap1 and ORC/Sir1 is to recruit Sir4/Sir2 and Sir3 to the silenced domain thus enabling these proteins to deacetylate and bind nucleosomes. The strength of the silencer is likely a function of the binding affinities of these proteins for silencer DNA in the context of nucleosomes and for the interactions of ORC with Sir1.

There are several possible explanations why strong housekeeping genes can be repressed by increasing levels of Sir1. The first point to make is that the binding affinity of Gal4-Sir1 for Gal4 binding sites at the synthetic silencer is unlikely to be the same as the binding affinity of Sir1 for ORC at the native silencer and therefore the dwell times of these two proteins at the silencer could be different thereby affecting the efficiency of silencing. In addition, increasing the number of Sir1 molecules at the silencer may lead to increased local concentrations of the Sir2, Sir3 and Sir4 proteins at the silencers. Loss of Sir1 can be compensated by overproduction of Sir3 and Sir4 ^108^ and recent analysis in cells mutated for Sir1 shows that a key function of silencer bound Sir1 is robust recruitment of the Sir proteins and their spread from the silencers^90^. The increase in Sir protein concentration at silencers is unlikely to alter the binding affinity of the Sir complexes for individual nucleosomes but the increased concentration could affect the search times required by the Sir proteins to find, deacetylate and bind unacetylated nucleosomes and this in turn would affect the ability of histone acetylases and chromatin remodellers to find, acetylate and mobilize nucleosomes thus altering the kinetic parameters of activation which in turn would alter the probability mass of Sir bound nucleosomes and silencing.

### Bistability Versus Monostability

Models have been proposed to help explain the stability of the silent state. Along with the silencers, a positive feedback loop and non-linearity (cooperativity), create a dispersed chromatin state where each nucleosome acts as a weak point silencer. This helps in the spreading of the Sir complex which then results in a three-dimensional mesh of protein-protein-nucleosome interactions leading to the formation of a highly stable silencing state^87,109-112^.

The stability of the active state that operates in opposition to stable silencing is a key feature undergirding eukaryotic differentiation and development. In *S. cerevisiae*, the bistable expression state arises under very specific conditions ^90^ and is dependent upon the structure and molecular characteristics of specific regulatory elements. Transcription activation is a multi-step process where each step is inefficient and probabilistic thus generating stochastic bursts of transcription. Activation is therefore heterogeneous because gene regulatory elements do not continuously engage the transcription machinery ^37,113,114^. The long-term stability of gene activation (both monostable activation and bistable expression) is likely determined by numerous biochemical factors including the number of binding sites for transcription factors in the regulatory regions of genes, the concentrations of transcription factors and the binding characteristics of these proteins in the context of chromatin modifications and remodelling. In the context of bistable expression states, one factor that likely helps the stable active state to persist is the observation that the establishment of the silent state is an inherently slower process and requires an extended period of transcription quiescence to form ^115^. This difference in the temporal kinetics of formation of the active and silent state allows the active state to persist and escape silencing. A gene that is repeatedly activated would thus allow the active state to be stably propagated leading to the formation of the bistable expression state.

Collectively, our data suggest that yeast silenced chromatin has developed to repress and stably silence transcription from weak constitutively active gene regulatory elements. Whether or not the same holds true for silencing in larger eukaryotes requires further analysis but consistent with this model are the observations that locus control regions in mammalian cells can overcome gene silencing (reviewed in ^116^). Furthermore, the demonstration that expressing a small number of mammalian pioneer transcription factors is sufficient to overcome facultative heterochromatin mediated silencing and cell differentiation ^117,118^ is also consistent with this model. It should also be noted that the boundaries between competing chromatin domains in all organisms are often occupied by housekeeping genes ^7,119–125^.

## Acknowledgements

We would like to thank Dr. Needhi Bhalla for the use of her wide-field fluorescence microscope and Dr. Susan Carpenter for the use of her Attune Cytometer. We would also like to thank Robert Shelansky for the gift of the *PP7-PHO5* coding region cassette.

We would like to thank Bailen Lawson and Caroline Olsmat for help with the analysis of images from the MFM microscope. We would like to thank Masaya Oki and Dave Clark for comments on the manuscript. The very early part of this work was funded by grants from the NIH to RTK (GM078068) and KW (T32-GM008646).

---

**Supplementary Figure 1.**
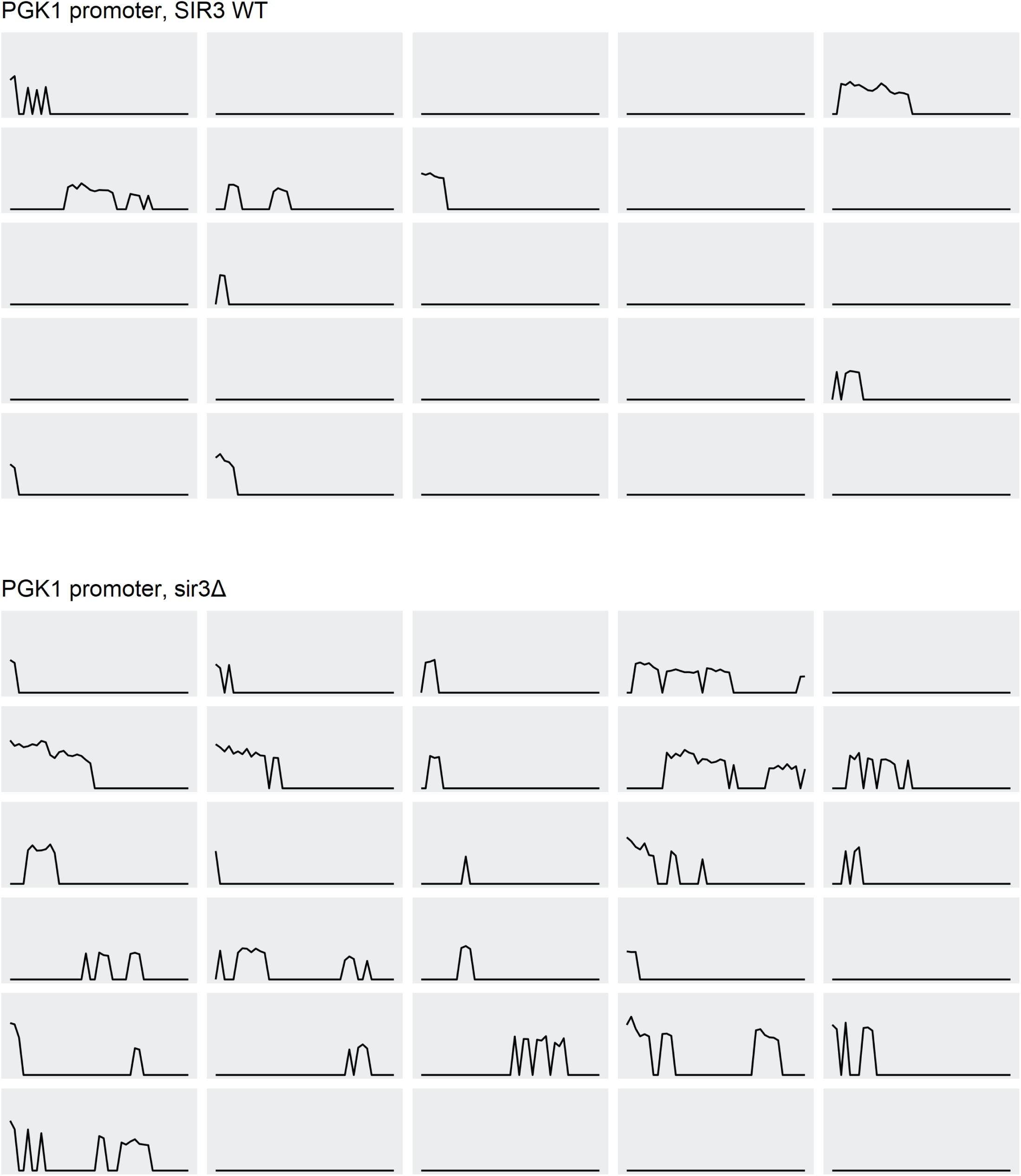
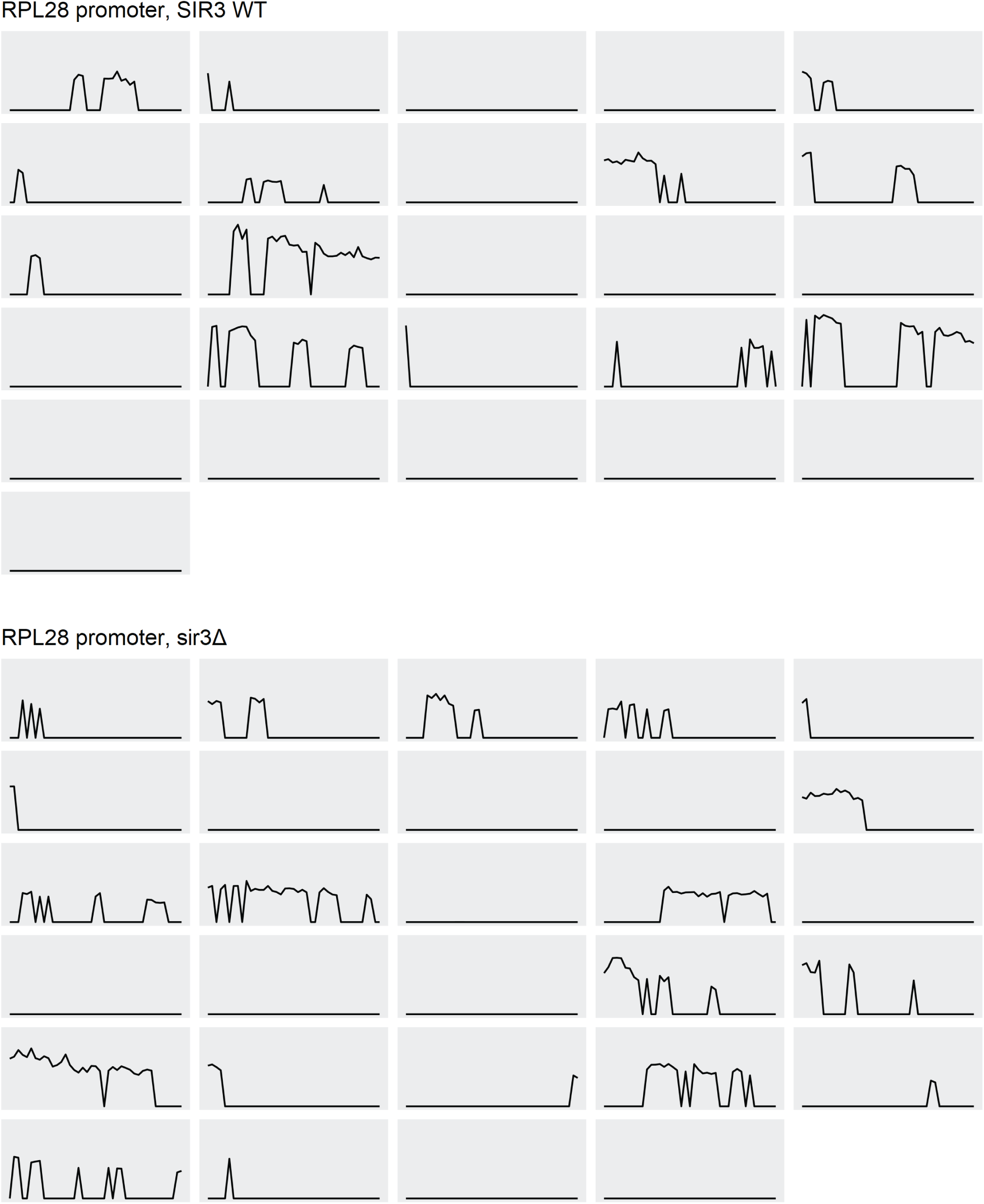
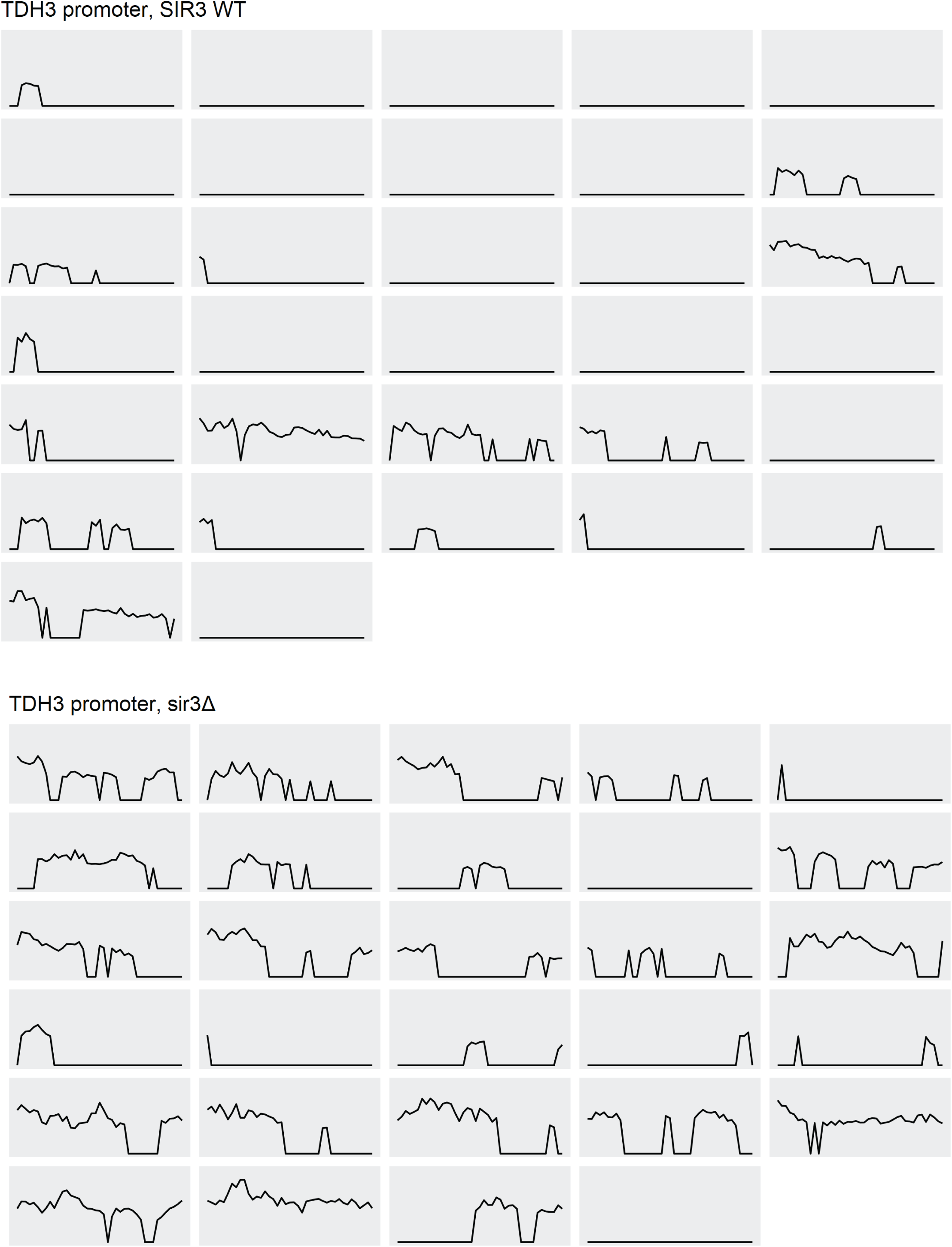
Fluorescent dot intensity traces of individual cells.

## Notes

### Competing Interest Statement

The authors have declared no competing interest.

